# A pretectal command system controls hunting behaviour

**DOI:** 10.1101/637215

**Authors:** Paride Antinucci, Mónica Folgueira, Isaac H. Bianco

## Abstract

For many species, hunting is an innate behaviour that is crucial for survival, yet the circuits that control predatory action sequences are poorly understood. We used larval zebrafish to identify a command system that controls hunting. By combining calcium imaging with a virtual hunting assay, we identified a discrete pretectal region that is selectively active when animals initiate hunting. Targeted genetic labelling allowed us to examine the function and morphology of individual cells and identify two classes of pretectal neuron that project to ipsilateral optic tectum or the contralateral tegmentum. Optogenetic stimulation of single neurons of either class was able to induce sustained hunting sequences, in the absence of prey. Furthermore, laser ablation of these neurons impaired prey-catching and prevented induction of hunting by optogenetic stimulation of the anterior-ventral tectum. In sum, we define a specific population of pretectal neurons that functions as a command system to drive predatory behaviour.

**Key findings:**

- Pretectal neurons are recruited during hunting initiation
- Optogenetic stimulation of single pretectal neurons can induce predatory behaviour
- Ablation of pretectal neurons impairs hunting
- Pretectal cells comprise a command system controlling hunting behaviour

## Introduction

In response to sensory information and internal states, animals select specific actions from a repertoire of options and produce adaptive behavioural programmes. Neuroethological studies in a variety of species have pinpointed brain regions, and identified neurons, that specifically promote particular behaviours ranging from the courtship songs of fruit flies (von Philipsborn *et al.*, 2011) to parental behaviours of mice (Kohl *et al.*, 2018). Identifying the neural circuits that control specific behaviours, as well as modulatory systems that influence if and how behaviours are performed, will shed light on the neural mechanisms of decision making, action selection and motor sequence generation.

Prey catching is an innate, complex behaviour that is crucial for survival (Sillar *et al.*, 2016). In various species, hunting responses can be evoked by prey-like stimuli, defined by specific conjunctions of sensory features (Ewert, 1997; Anjum *et al.*, 2006; Bianco and Engert, 2015), and predatory behaviour is modulated by internal state variables including associative learning and feeding drive (Ewert *et al.*, 2001; Jordi *et al.*, 2015). Several brain regions show activity during hunting and are expected to subserve neural functions including prey detection and localisation, control of pursuit, capture and consummatory actions, and motivation [*e.g.* Comoli *et al.* (2005)]. Although electrical stimulation of brain regions, including the optic tectum, can evoke hunting actions (Ewert, 1970; Bels *et al.*, 2012) and recent studies in rodents have identified circuits that motivate predatory behaviour (Han *et al.*, 2017; Li *et al.*, 2018; Park *et al.*, 2018), neurons that command vertebrate hunting have yet to be identified.

In this study, we sought to identify a command system for control of predatory behaviour, using larval zebrafish as a vertebrate model system. Command systems comprise interneurons that are activated in association with a specific behaviour and whose activation is able to induce that behaviour (Kupfermann and Weiss, 1978; Yoshihara and Yoshihara, 2018). In contrast to modulatory circuits, the presence of the releasing stimulus should not be required for experimental activation of command neurons to induce the behavioural response. In larval zebrafish, hunting is an innate, visually guided behaviour, which involves a sequence of specialised oculomotor and locomotor actions. A defining characteristic is that larvae initiate hunting by rapidly converging their eyes, which substantially increases their binocular visual field (Bianco *et al.*, 2011). A high vergence angle is maintained during prey pursuit, which entails a sequence of discrete orienting turns and approach swims, which culminate in binocular fixation of prey followed by a kinematically distinct capture swim (Borla *et al.*, 2002; McElligott and O’Malley D, 2005; Bianco *et al.*, 2011; Patterson *et al.*, 2013; Trivedi and Bollmann, 2013; Marques *et al.*, 2018). Neural activity associated with prey-like visual cues has been detected in the axon terminals of a specific class of retinal ganglion cell (RGC), which terminate in retinal arborization field 7 (AF7) in the pretectum (Semmelhack *et al.*, 2014). Prey-responsive pretectal cells have also been described (Muto *et al.*, 2017) as well as highly prey-selective feature-analysing neurons in the optic tectum (OT) that display non-linear mixed selectivity for conjunctions of visual features (Bianco and Engert, 2015). Premotor activity in localised tectal assemblies immediately precedes execution of hunting responses (Bianco and Engert, 2015) and optogenetic stimulation of the anterior-ventral tectal region can induce hunting-like behaviour (Fajardo *et al.*, 2013). Finally, ablation of RGC input to either AF7 or OT substantially impairs hunting (Gahtan *et al.*, 2005; Semmelhack *et al.*, 2014).

Based on this evidence, we hypothesised that neural circuits controlling the induction of hunting might be located in the vicinity of AF7 or OT. Our experimental requirements for identifying neurons that fulfil the criteria of a command system were (1) that they display neural activity specifically related to execution of hunting behaviour, rather than visual detection of prey, and (2) that direct stimulation of such neurons would induce naturalistic predatory behaviour in the absence of prey. First, we used 2-photon calcium imaging paired with a virtual hunting assay and identified a high density of neurons in the AF7-pretectal region that were specifically recruited when larvae initiated hunting behaviour. We identified a transgenic line that labelled these neurons and found two morphological classes: One projects ipsilaterally to the optic tectum and the second extends long-range projections to midbrain oculomotor nuclei, the nucleus of the medial longitudinal fasciculus and the contralateral hindbrain tegmentum. Remarkably, optogenetic stimulation of single pretectal neurons evoked hunting-like behaviour in the absence of prey. Pretectal projection neurons of either class could evoke hunting routines with naturalistic oculomotor and locomotor kinematics but opposite directional biases. Finally, laser-ablation of the pretectal population impaired hunting of live prey. In sum, we propose that a specific population of pretectal neurons comprises a command system that functions downstream of prey perception to control execution of predatory behaviour.

## Results

### Pretectal neurons are recruited during hunting initiation

To identify neurons with activity related to prey perception and/or initiation of hunting behaviour, we performed 2-photon calcium imaging while larval zebrafish engaged in a virtual hunting assay (Figure 1A) (Bianco and Engert, 2015). Transgenic *elavl3:H2B-GCaMP6s;atoh7:gapRFP* larvae (6–7 dpf, N = 8) were partially restrained in agarose gel, but with their eyes and tail free to move, and were presented with a range of visual cues including small moving prey-like spots, which evoke naturalistic hunting responses (Figure 1B) (Bianco *et al.*, 2011; Bianco and Engert, 2015). We imaged a volume that encompassed the majority of the primary retinorecipient sites [arborization fields (AFs) 2–10] as well as surrounding brain regions including pretectum and OT (310 × 310 × 100 *µ*m volume; Figure 1C and Video 1). Eye and tail kinematics were tracked online, allowing automated detection of hunting responses, which are defined by saccadic convergence of the eyes – a unique oculomotor behaviour executed exclusively at hunting initiation (Figure 1D) (Bianco *et al.*, 2011; Patterson *et al.*, 2013; Trivedi and Bollmann, 2013; Bianco and Engert, 2015). Larvae preferentially responded to small, dark, moving spots and hunting was initiated most frequently once the stimulus had crossed the midline axis and was moving in a nose-tail direction (Figure 1E,F) (Bianco and Engert, 2015).

**Figure 1.**
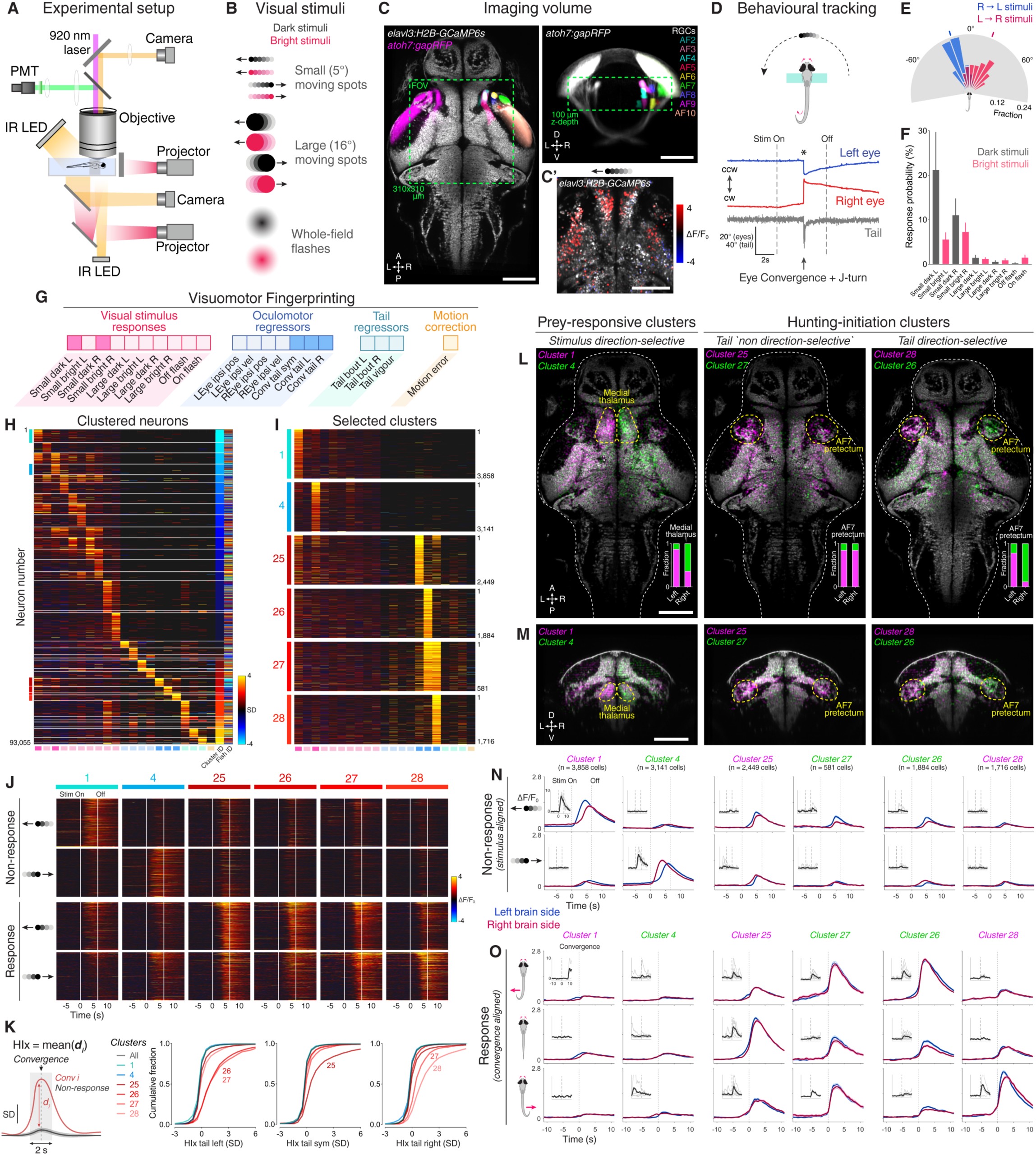
AF7-pretectal neurons are recruited at onset of hunting behaviour. **A** 2-photon GCaMP imaging combined with behavioural tracking during virtual hunting behaviour (see Materials and Methods). **B** Schematic of visual stimuli. **C** *elavl3:H2B-GCaMP6s;atoh7:gapRFP* reference brain showing imaging volume (green), which encompassed most retinal arborisation fields (AF2–10). RFP has been pseudo-coloured to demarcate specific AFs on the right hemisphere. **C’** Example of neuronal activity (ΔF/F_0_) within one focal plane in response to a dark, leftwards moving prey-like spot (mean activity over 8 presentations) overlaid onto anatomical image (grey). **D** Example of behavioural tracking data indicating hunting initiation (eye convergence and leftwards J-turn) in response to a dark, leftwards moving prey-like spot. Asterisk indicates time of convergent saccade. cw, clockwise; ccw, counter-clockwise. **E** Distribution of spot locations at time of convergent saccade. Ticks indicates median location for leftwards (blue, –18.13°, n = 162 events in 8 fish) and rightwards (red, 22.10°, n = 122 events) moving spots. **F** Hunting response probability (mean + SEM, N = 8 fish) across visual stimuli. **G** Schematic of the visuomotor vector (VMV) generated for each neuron (see Supplementary File 1 and Materials and Methods for detailed description). **H** VMVs of all clustered neurons (n = 93,055 neurons from 8 fish). White lines indicate cluster boundaries. For each cluster, neurons are ordered according to decreasing correlation with the cluster mean. Coloured lines on the left along the y-axis indicate hunting-related clusters with anatomical maps and ΔF/F_0_ responses shown in 1L–1O (prey-responsive clusters in blue, hunting-initiation clusters in red). **I** VMVs of selected clusters (1, 4, 25–28). Number of cells in each shown on right. **J** Stimulus-aligned activity during non-response (top) and response (bottom) trials for neurons in selected clusters (indicated top). **K** Hunting Index (HIx) for selected clusters. Left shows schematic indicating how HIx is computed and right shows distribution of HIx scores for selected clusters. **L** Anatomical maps of prey-responsive clusters (left) and hunting-initiation clusters (middle and right). Images show dorsal views of intensity sum projections of all neuronal masks in each cluster after registration to the *elavl3:H2B-GCaMP6s* reference brain (grey). Insets show fraction of neurons in left and right AF7-pretectum or medial thalamus belonging to specified clusters. **M** Ventro-dorsal cross-section views of anatomical maps. **N** Visual stimulus-aligned activity during non-response trials for prey-responsive clusters (left) and hunting-initiation clusters (middle and right; mean ± SEM). Traces are colour-coded according to anatomical laterality (blue, left; red, right). Insets show single-trial responses for a single example cell from each cluster (mean as thick line). **O** Eye convergence-aligned activity for convergences associated with leftwards (top), rightwards (bottom), or symmetrical/no tail movements (middle). Activity during both spontaneous and visually evoked convergences was used. Scale bars, 100 *µ*m. A, anterior; D, dorsal; L, leftwards; P, posterior; R, rightwards; V, ventral; Sym, symmetric. See also Figure 1–supplement 1–3, and Video 1.

To define groups of neurons with consistent functional properties related to the first stages of hunting behaviour, we first computed, for every cell, a visuomotor vector (VMV) that quantified its sensory and motor-related activity (Figure 1G and Materials and Methods). Each VMV described (a) mean GCaMP fluorescence responses to each of the ten visual cues during non-response trials, in which larvae did not release hunting behaviour, and (b) coefficients from multilinear regression of fluorescent timeseries data on a set of motor predictors (derived from eye and tail kinematics including convergent saccades; see Supplementary File 1). Next, we used an unsupervised clustering procedure to identify consistent sensorimotor tuning profiles. A correlation-based agglomerative hierarchical clustering algorithm performed initial clustering of VMVs from cells with either high visually evoked activity or that were well modelled in terms of motor variables (see Materials and Methods). The centroids of the resultant clusters defined a set of functional archetypes and subsequently, all remaining neurons were assigned to the cluster with the closest centroid within a threshold distance limit (Pearson’s *r* ≥ 0.7, VMVs below threshold remained unassigned). This approach allowed us to classify more than 50% of imaged neurons (93,055 out of 181,123 cells) into 36 clusters with homogenous functional properties (Figure 1H and Figure 1–supplement 1).

This analysis identified neurons that were recruited during hunting initiation. Specifically, four clusters showed activity highly correlated with eye convergence (clusters 25–28; Figure 1I) yet exhibited little activity in response to visual cues (including prey-like moving spots; Figure 1J and Figure 1–supplement 2A). In two of these clusters, activity was selective for the direction of tail movements that often occur concomitantly with eye convergence during hunting initiation. Thus, cluster 26 was selective for convergences associated with leftwards turns and cluster 28 was tuned to rightwards hunting responses. By contrast, the other two clusters did not show selectivity for the direction of tail movements during hunting initiation (cluster 25, associated with symmetrical/no tail movement and 27, responsive to motion in either direction). Other functional clusters comprised visually responsive neurons that were selectively activated by small prey-like moving spots (’prey-responsive’ clusters 1–6; Figure 1H,I and Figure 1–supplement 2A), but displayed little motor-related activity.

We computed a ‘hunting index’ (HIx) for individual neurons as a direct means to distinguish neural activity associated with hunting initiation from ‘sensory’ activity evoked by prey-like visual stimuli. Briefly, for each hunting response, GCaMP fluorescence in a time window (±1 s) surrounding the convergent saccade was compared to activity at the same time in non-response trials during which the same visual stimulus was presented (Figure 1K, left). The mean of these difference measures across all response events represents the HIx score for the cell and quantifies neural activity attributable to hunting initiation while accounting for any visually evoked response. To account for directional tuning, we separately computed HIx for hunting responses paired with leftwards, rightwards or symmetrical/no tail movements. This analysis revealed that neurons in clusters 25–28 showed considerably higher HIx scores than other cells, including those in prey-responsive clusters (1 and 4, Figure 1K). Moreover, tail directional preferences were consistent with those determined by regression modelling. Overall, our functional analyses identified four clusters of neurons with activity specifically associated with the specialised motor outputs that characterise initiation of hunting behaviour and showed little activity in response to prey-like visual cues. Thus, we will refer to these as ‘hunting-initiation’ clusters.

Neurons within functionally defined clusters showed distinct anatomical distributions (Figure 1–supplement 3). We showed this by registering volumetric imaging data to a reference brain atlas (‘ZBB’ and a high-resolution *elavl3:H2B-GCaMP6s* volume, see Materials and Methods and Video 1) (Marquart *et al.*, 2015; Marquart *et al.*, 2017). A high density of neurons belonging to hunting-initiation clusters was found in pretectal regions in the immediate vicinity of AF7 (AF7-pretectum), just anterior to the rostral pole of the optic tectum (Figure 1L,M and Figure 1–supplement 2B,C). Neurons selective for the direction of hunting-related tail motion (clusters 26 and 28) showed lateralised, mirror-symmetric distributions, with a larger fraction of cells located on the side of the brain contralateral to the direction of preferred tail movement (Figure 1L, right panel). Specifically, cluster 26, which is tuned to eye convergences associated with leftwards tail movement, had a higher density of cells in the right AF7-pretectum, and vice versa for cluster 28. Hunting-initiation clusters that were agnostic to tail direction (clusters 25 and 27) showed largely symmetric anatomical distributions (Figure 1L, middle panel). Neurons belonging to prey-responsive clusters were also found in AF7-pretectum as well as in the medial thalamus, where direction-selective neurons showed highly lateralised distributions (Figure 1L,M left panels and Figure 1– supplement 3).

To confirm the response properties indicated by the VMV representations and HIx scores and further examine visuomotor tuning, we computed visual stimulus-aligned and convergence-aligned activity profiles for left and right hemisphere neurons in prey-responsive and hunting-initiation clusters (Figure 1N,O). This confirmed that prey-responsive neurons in clusters 1 and 4 showed direction-selective activity in response to small dark moving spots, but minimal activity associated with eye convergence (Figure 1N,O, left columns). On the other hand, hunting-initiation neurons (clusters 25–28) showed weak visual responses – as shown by moving spot-triggered activity during non-response trials – but substantial activity triggered on hunting initiation. For clusters 26 and 28, neurons showed stronger activation when convergent saccades were paired with left and right-sided turns, respectively (Figure 1N,O, right columns).

In summary, we identified populations of neurons in AF7-pretectum that are recruited in association with two distinct components of hunting – visual responses to prey and initiation of predatory behaviour. We subsequently examined whether cells with hunting-initiation activity are directly involved in inducing hunting behaviour.

### Pretectal neurons labelled by *KalTA4u508* with hunting-initiation activity

To characterise the connectivity and function of AF7-pretectal neurons with hunting-initiation activity, we inspected the expression patterns of a range of transgenic driver lines and identified a transgene, *KalTA4u508*, which preferentially labels neurons in the AF7-pretectal region (Figure 2A–C). Anatomical registration of *KalTA4u508;UAS:mCherry* volumetric data to the reference atlas revealed a high density of labelled somata in AF7-pretectum, overlapping with the locations of hunting-initiation clusters (Figure 2A,B,I). We generated *KalTA4u508;UAS:RFP;atoh7:GFP* larvae to visualise GFP-labelled RGC axon terminals in AF7, and observed that a subset of *KalTA4u508-*expressing neurons extend dendritic arbours that directly juxtapose RGC terminals (N = 4 fish; Figure 2C,D). In brain sections from adult (3 month old) *KalTA4u508;UAS:GCaMP6f;atoh7:gapRFP* fish, labelled neurons in the pretectum were very sparse and could be identified only in the accessory pretectal nucleus (APN) (n = 8 somata from N = 4 fish; Figure 2E). This suggests that at least a subset of *KalTA4u508-*expressing neurons reside in a region of the larval AF7-pretectum corresponding to the adult APN.

**Figure 2.**
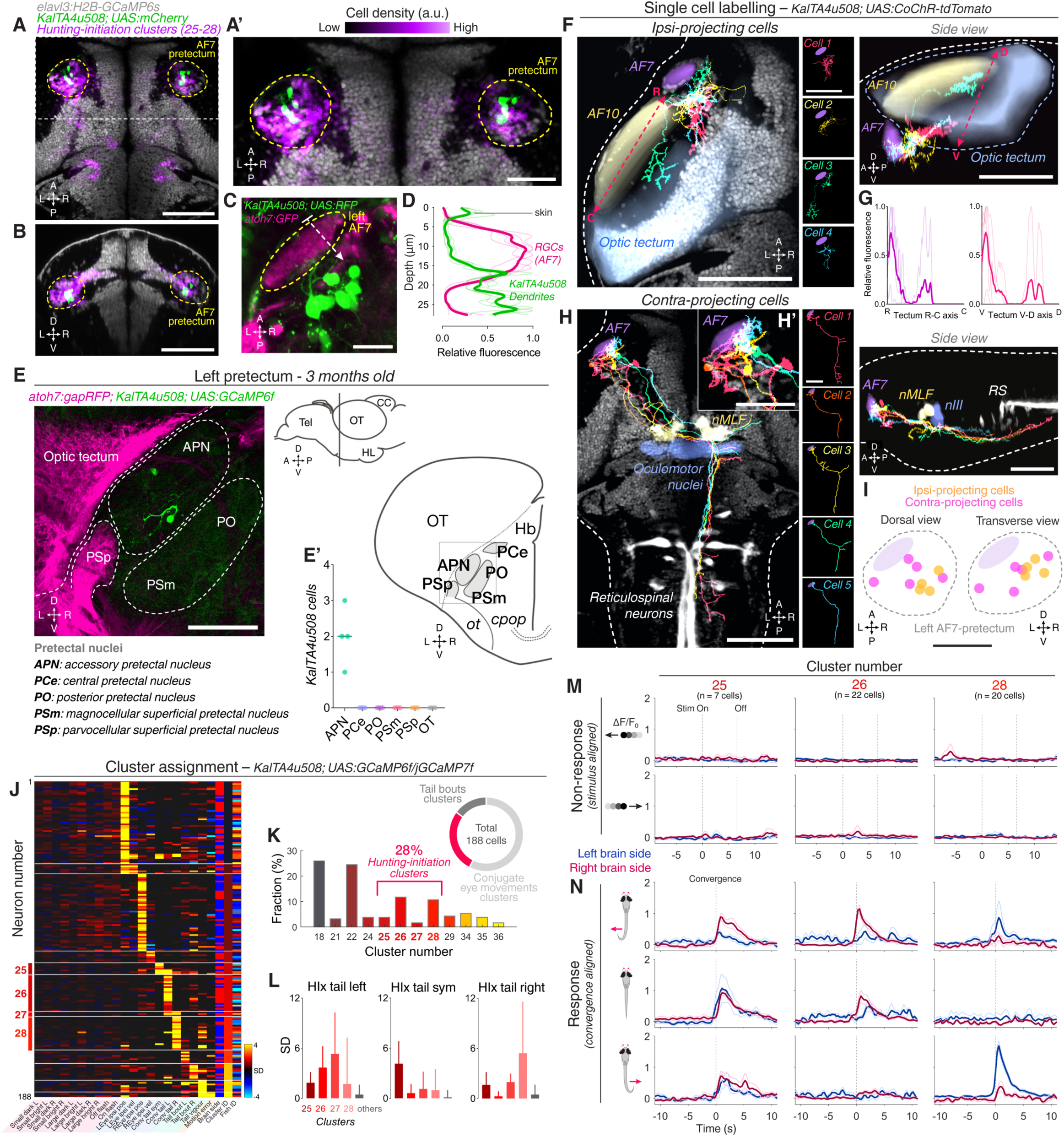
Pretectal neurons labelled by *KalTA4u508* are active during hunting initiation. **A** Dorsal view of *KalTA4u508;UAS:mCherry* expression at 6 dpf (green) registered to the *elavl3:H2B-GCaMP6s* reference brain (grey). Neurons of all four hunting-initiation clusters combined are shown in purple, colour-coded according to local cell density (clusters 25–28, n = 6,630 cells from 8 fish). AF7-pretectum is indicated in yellow and the region is enlarged in **A’**. **B** Ventro-dorsal cross-section of data in **A**. **C** Left AF7-pretectum in a 6 dpf *KalTA4u508;UAS:RFP;atoh7:GFP* larva (dorsal view, maximum-intensity projections, 10 planes, 10 *µ*m depth). **D** Dendritic stratification of *KalTA4u508* neurons (green) relative to RGC axons (magenta) in AF7. Y-axis indicates distance from the skin in *µ*m (dashed white arrow in C). Mean and individual stratification patterns are reported (N = 4 fish). **E** *KalTA4u508* neurons in pretectum of a 3 month-old *KalTA4u508;UAS:GCaMP6f;atoh7:gapRFP* fish. Pretectal and tectal regions in the left hemisphere are shown. Schematic indicates location of micrograph and pretectal nuclei (transverse plane). Number of *KalTA4u508* cells in each pretectal nucleus are reported in **E’** (N = 4 fish). APN, accessory pretectal nucleus; CC, cerebellar corpus; *cpop*, postoptic commissure; Hb, habenula; HL, hypothalamic lobe; OT, optic tectum; *ot*, optic tract; PCe, central pretectal nucleus; PO, posterior pretectal nucleus; PSm, magnocellular superficial pretectal nucleus, PSp, parvocellular superficial pretectal nucleus; Tel, telencephalon. **F** Tracings of individually labelled *KalTA4u508* projection neurons that innervate the ipsilateral tectum (‘ipsi-projecting’ cells, n = 4) in 6–7 dpf *KalTA4u508;UAS:CoChR-tdTomato;elavl3:H2B-GCaMP6s* larvae registered to the *elavl3:H2B-GCaMP6s* reference brain (grey). Selected anatomical regions from the ZBB brain atlas are overlaid. To enable morphological comparisons, all traced neurons are shown in the left hemisphere. **G** Fluorescence profiles of neurites of ipsi-projecting *KalTA4u508* cells along the rostro-caudal (R-C, left) and ventral-dorsal (V-D, right) axes of the optic tectum (dashed red arrows in **F**). Mean and individual profiles are reported (n = 4 cells). **H** Tracings of *KalTA4u508* projection neurons innervating the contralateral hindbrain (‘contra-projecting’ cells, n = 5). Dendritic arbours adjacent to AF7 are enlarged in inset **H’**. nMLF, nucleus of the medial longitudinal fasciculus; RS, reticulospinal system. **I** Soma location of ipsi- and contra-projecting *KalTA4u508* cells in AF7-pretectum. **J** VMVs of *KalTA4u508* neurons with assigned cluster identities (n = 188 neurons from 30 fish). Cell location (blue for left hemisphere, red for right) is reported by the ‘Brain side’ column. **K** Fraction of assigned *KalTA4u508* neurons in each cluster. **L** Hunting Index (HIx) scores for *KalTA4u508* neurons in different clusters (mean + SD). **M** Visual stimulus-aligned responses of *KalTA4u508* neurons during non-response trials (mean ± SEM). Traces are colour-coded according to anatomical laterality (blue for left hemisphere, red for right). **N** Eye convergence-aligned neuronal responses. Activity during both spontaneous and visually evoked convergences was used. Scale bars, 100 *µ*m, except **A’, H’, I**, 50 *µ*m, and **C**, 20 *µ*m. A, anterior; C, caudal; D, dorsal; L, left; P, posterior; R, right (rostral in **G**); V, ventral; Sym, symmetric; Stim, stimulus. See also Figure 2–supplement 1.

To reveal the morphology of *KalTA4u508* neurons, we used a transient expression strategy to label individual cells by microinjection of a *UAS:CoChR-tdTomato* DNA construct into one-cell stage *KalTA4u508;elavl3:H2B-GCaMP6s* embryos. High-contrast membrane labelling by CoChR-tdTomato facilitated morphological reconstruction of single neurons (at 6–7 dpf) and tracings were registered to the brain atlas using the H2B-GCaMP6s channel (Figure 2F,H).

We identified three morphological classes of pretectal neuron labelled by *KalTA4u508*. One class projects to the ipsilateral optic tectum (Figure 2F), elaborating axon terminal arbours preferentially in the most anterior-ventral aspect of OT (n = 4 cells from 4 fish; Figure 2G). The second class makes descending projections to the midbrain and hindbrain tegmentum. Axons decussate near the nucleus of the medial longitudinal fasciculus (nMLF) and the oculomotor nucleus (nIII) before extending caudally into the contralateral ventral hindbrain. Axon collaterals could be observed bilaterally in the vicinity of nIII/nMLF and proximal to ventral reticulospinal neurons in the contralateral hindbrain (n = 5 cells from 5 fish; Figure 2H). This class of projection neuron extended dendrites within a neuropil region that includes AF7 (Figure 2H’), a feature not observed in the other two classes. The third class was characterised by ipsilateral axonal projections to the medial region of the corpus cerebellum (n = 2 cells from 2 fish; Figure 2–supplement 1A). Neurite tracing using photoactivatable GFP confirmed a pretectal projection to the cerebellum (Figure 2–supplement 1C) as well as to nIII/nMLF and contralateral ventral hindbrain (Figure 2–supplement 1B). This latter projection pattern is compatible with data on APN efferent projections in adult zebrafish (Yanez *et al.*, 2018). In combination with our adult expression data (Figure 2E), we conclude that the subset of *KalTA4u508* pretectal neurons projecting to contralateral hindbrain belong to the larval APN.

Next, we asked whether *KalTA4u508* pretectal neurons are responsive to prey-like stimuli and/or are recruited during hunting initiation. We performed 2-photon calcium imaging in *KalTA4u508;UAS:GCaMP6f* or *KalTA4u508;UAS:GCaMP7f* transgenic larvae during the virtual hunting assay (6–7 dpf, N = 30 fish). Notably, *KalTA4u508* pretectal neurons exhibited negligible activity in response to visual stimuli (Figure 2M and Figure 2– supplement 1E). Visuomotor vectors were generated for individual cells allowing ∼51% to be assigned cluster identities based on the functional archetypes established previously using pan-neuronal imaging (correlation threshold = 0.7, n = 188 out of 369 cells).

Of the *KalTA4u508* cells assigned functional identities, 28% were associated with hunting-initiation clusters (clusters 25–28; 52/188 cells; Figure 2J,K). The remaining neurons were assigned to either conjugate eye movement clusters (57%) or tail movement clusters (15%) and, in line with the absence of visual sensory responses, no cells were assigned to prey-responsive clusters. Of the hunting-initiation neurons, most *KalTA4u508* cells were associated with functional clusters 26 and 28 which show preference for tail direction coincident with eye convergence (Figure 2K,N). As before, *KalTA4u508* pretectal neurons in these two clusters were predominantly located contralateral to the direction of preferred tail movement (73% and 80% contralateral for cluster identities 26 and 28, respectively; Figure 2J,N). *KalTA4u508* cells assigned to hunting-initiation clusters had higher HIx scores than those assigned to other clusters, supporting the hunting-response specificity of their activity (Figure 2L,N).

In summary, *KalTA4u508* provides genetic access to a subset of AF7-pretectal neurons that are selectively active during initiation of hunting behaviour.

### Optogenetic activation of single *KalTA4u508* pretectal neurons induces hunting

To test whether *KalTA4u508* pretectal cells are capable of inducing predatory behaviour, we optogenetically stimulated single neurons while using high-speed tracking to monitor free-swimming behaviour (Figure 3A). To do this, we used the same larvae described above in which single *KalTA4u508* pretectal cells expressed the optogenetic actuator CoChR-tdTomato (Figure 3B) (Klapoetke *et al.*, 2014). Experiments consisted of repeated trials, each of 8 s duration, in which larvae (6–7 dpf, N = 70) were exposed to 7 s blue light stimulation (470 nm, 0.44 mW/mm^2^), interleaved with trials with no stimulation.

**Figure 3.**
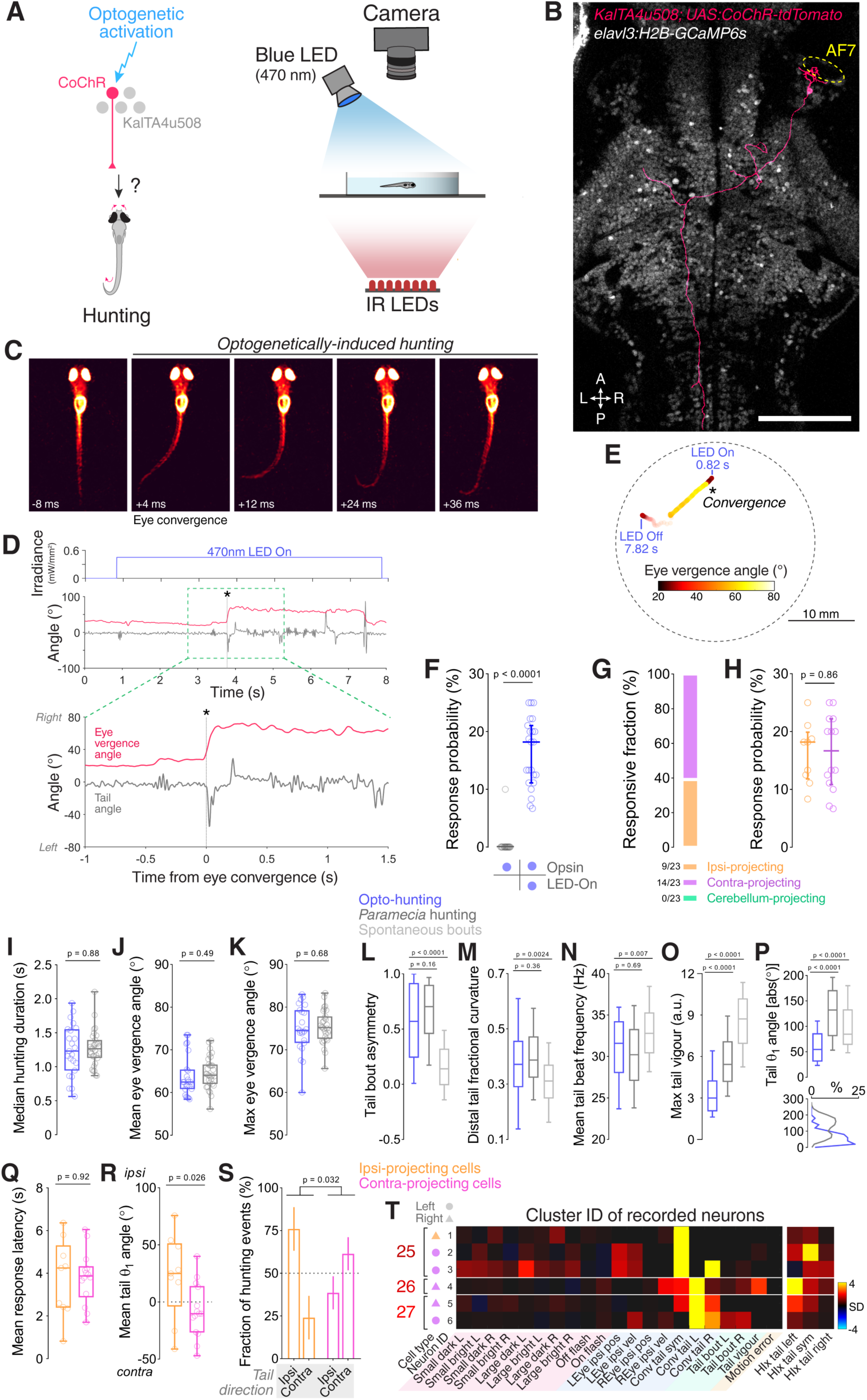
Optogenetic stimulation of single *KalTA4u508* pretectal neurons induces hunting. **A** Optogenetic stimulation of single neurons paired with behavioural tracking. **B** A single *KalTA4u508* neuron in a 7 dpf *KalTA4u508;elavl3:H2B-GCaMP6s* larva that was injected at the one-cell stage with *UAS:CoChR-tdTomato* DNA. This ‘contra-projecting’ neuron is ‘Cell 3’ in Figure 2H. A, anterior; L, left; P, posterior; R, right. Scale bar, 100 *µ*m. **C** Example frames from an optogenetically induced hunting event. Labels indicate time relative to saccadic eye convergence, which marks hunting initiation (t = 0 ms). **D** Behavioural tracking of tail angle (grey) and ocular vergence angle (red) during an optogenetically induced hunting event (ipsi-projecting cell located in the left AF7-pretectum, this neuron is ‘Cell 4’ in Figure 2F; see also Video 2). Asterisk indicates time of convergent saccade. **E** Larval location colour-coded by vergence angle during the example hunting event in **D** (see also Video 2). **F** Hunting response probability in LED-On versus non-stimulation trials for larvae that performed at least one eye convergence during optogenetic stimulation (N = 23 fish). **G** Morphological identity of *KalTA4u508* neurons that elicited hunting upon optogenetic stimulation. Numbers of responsive larvae are reported at the bottom (N = 23 fish). **H** Hunting response probability upon optogenetic stimulation of ipsi-projecting (orange, n = 9 cells) and contra-projecting neurons (magenta, n = 14 cells). **I–P** Comparison of behavioural kinematics between optogenetically induced hunting events (blue, N = 23 fish) and *Paramecia* hunting (dark grey, N = 31 fish). Tail kinematics for non-hunting swim bouts were recorded from larvae that were monitored during *Paramecia* hunting (light grey). In **L–P**, data from all bouts are plotted, whereas in **I–K** the median, mean or maximum for each larva is reported. **Q–S** Behavioural kinematics of hunting events induced by stimulation of ipsi-projecting *KalTA4u508* neurons (orange, n = 9 cells) or contra-projecting neurons (magenta, n = 14 cells). **T** VMVs and cluster identity of *KalTA4u508* neurons that induced hunting upon optogenetic stimulation and subsequently underwent calcium imaging (n = 6 cells from 6 larvae). Symbols on the left indicate projection cell class and left/right location, and HIx scores are shown on right. See also Figure 3–supplement 1 and Video 2.

Strikingly, we found that optogenetic stimulation of individual *KalTA4u508* pretectal neurons could induce sustained, hunting-like behavioural routines. The fraction of *KalTA4u508* pretectal neurons that induced hunting (32%, 23 out of 70 cells) was similar to the proportion that were assigned to hunting-initiation clusters (28%; Figure 2K). As in naturalistic hunting, optogenetically induced hunting routines were initiated with convergent saccades accompanied by lateralised swim bouts (Figure 3C) and often continued for several seconds (Figure 3D,E, and Video 2). Hunting-like responses were entirely dependent on blue light stimulation. For responsive fish we observed 18.1% median response probability in LED-On trials vs. 0% in LED-Off trials (p < 0.0001, N = 23 fish; Figure 3F). Furthermore, control experiments demonstrated that opsin-negative animals do not produce hunting behaviours in response to blue light stimulation alone (Figure 4–supplement 1I,J). By examining single-cell morphology, we found that the *KalTA4u508* cells that could evoke hunting behaviour belonged to the projection classes that innervated the ipsilateral optic tectum (9/23 cells, hereafter abbreviated ‘ipsi-projecting’) or that belong to the presumptive APN and connect to the contralateral tegmentum (14/23 cells, ‘contra-projecting’; Figure 3G,H).

**Figure 4.**
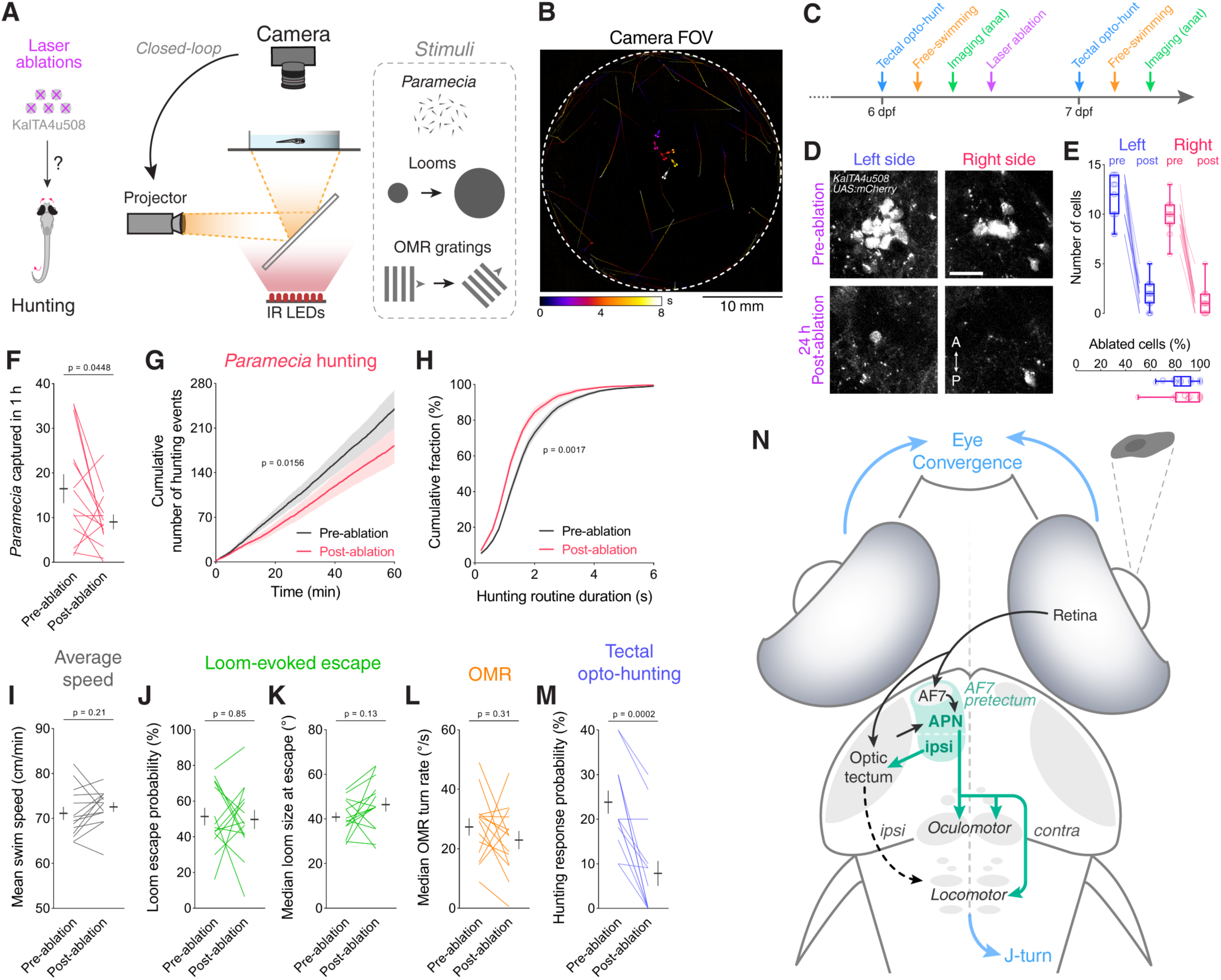
Ablation of *KalTA4u508* pretectal neurons impairs hunting. **A** Laser ablation of *KalTA4u508* pretectal neurons and assessment of visuomotor behaviours. **B** Time-projection of larval behaviour (duration 8 s) showing trajectories of *Paramecia* and larval zebrafish swimming in the arena. **C** Time course of behavioural tests, ablation and brain imaging. **D** Pretectal neurons before (top, 6 dpf) and 24 h after (bottom, 7 dpf) bilateral laser ablations in a *KalTA4u508;UAS:mCherry;elavl3:itTA;Ptet:ChR2-YFP* larva. Images show maximum-intensity projections (red channel, 10 planes, 10 *µ*m depth). A, anterior; P, posterior. Scale bar, 20 *µ*m. **E** Quantification of cell ablation in left (blue) and right (red) AF7-pretectum (N = 14 larvae). **F–H** Assessment of hunting performance before and after bilateral ablation of *KalTA4u508* neurons (N = 14 larvae). Mean ± SEM is reported for each condition. **I** Average swim speed before and after ablations. **J–K** Loom-evoked escape behaviour before and after ablations. **L** OMR behaviour before and after ablations, quantified by rate of reorientation to a grating drifting at 90° with respect to the fish in a fixed egocentric reference frame. **M** Hunting response probability for optogenetic stimulation of *KalTA4u508;UAS:mCherry;elavl3:itTA;Ptet:ChR2-YFP* larvae before and after ablation of *KalTA4u508* pretectal neurons (N = 14 larvae). **N** Model of the neural circuit controlling hunting initiation. Two classes of AF7-pretectal projection neuron are capable of inducing hunting behaviour. Contralaterally projecting APN neurons are likely to induce hunting by recruiting activity in oculomotor and locomotor pattern generating circuits in the mid/hindbrain tegmentum. Ipsilaterally projecting AF7-pretectal neurons may recruit ipsilateral tectofugal pathways (Helmbrecht *et al.*, 2018). Activation of anterior-ventral tectum requires AF7-pretectal neurons to induce hunting and likely operates via the contralaterally projecting APN population because unilateral avOT stimulation produces contralaterally directed responses (Fajardo *et al.*, 2013). APN, accessory pretectal nucleus. See also Figure 4–supplement 1,2, and Video 3.

How closely do the hunting-like routines induced by optogenetic stimulation of *KalTA4u508* pretectal neurons compare to hunting behaviour? To address this question, we compared a variety of oculomotor and locomotor kinematics between optogenetically induced hunting routines versus hunting of live *Paramecia*. We observed no difference in mean or maximum ocular vergence angles between the two types of routine, and the duration of hunting sequences, defined by the period of elevated ocular vergence, was equivalent between natural and optogenetically evoked behaviour (Figure 3I–K). We analysed kinematic features of swim bouts associated with the convergent saccade that initiates hunting, focussing on features that distinguish hunting swims from swim bouts used during spontaneous exploratory behaviour. This revealed a high degree of kinematic similarity between natural hunting bouts and optogenetically induced hunting bouts. In both cases, swim bouts contained a highly lateralised sequence of half-beats (‘bout asymmetry’; Figure 3L, see Materials and Methods for details), a large fraction of curvature was localised to the distal segments of the tail (Figure 3M) and bouts displayed low tail beat frequency (Figure 3N). Notably, all such parameters were significantly different as compared to spontaneous swims. Optogenetically induced hunting bouts showed a reduction in average vigour and theta-1 angles (maximum tail angle during the first half beat) compared to bouts during *Paramecia* hunting (Figure 3O,P). However, the latter displayed a bimodal distribution of theta-1 and optogenetically induced hunting bouts overlapped with the lower amplitude component of this distribution (Figure 3P, bottom). In sum, our data indicate that optogenetic stimulation of single *KalTA4u508* pretectal neurons can induce naturalistic hunting-like behaviour.

Next, we compared hunting routines induced by optogenetic activation of ipsi-versus contra-projecting *KalTA4u508* neurons. We did not observe differences in response latency (Figure 3Q), sequence duration (Figure 3–supplement 1A) or oculomotor or tail kinematics (Figure 3–supplement 1B–F). However, the laterality of evoked hunting responses differed between the two projection classes (Figure 3R,S): Stimulation of ipsi-projecting *KalTA4u508* pretectal neurons most frequently induced hunting in which the first swim bout was oriented in the ipsilateral direction (*i.e.* a left pretectal neuron evoked leftward turning). By contrast, contra-projecting neurons most frequently induced contralaterally directed hunting responses, as might be expected from their axonal projections to contralateral tegmentum.

To directly establish whether the *KalTA4u508* pretectal neurons than can drive predatory behaviour are the same cells that are recruited during visually evoked hunting, we combined optogenetic stimulation and functional calcium imaging of single neurons. To achieve this, we first established that optogenetic stimulation of a given *KalTA4u508* pretectal neuron could induce hunting and then tethered the larva in agarose and performed calcium imaging of H2B-GCaMP6s, expressed in the nucleus of the same neuron, while the animal engaged in the virtual hunting assay. Visuomotor fingerprinting and cluster assignment of these neurons showed that they all belonged to hunting-initiation clusters (clusters 25–27) and had high HIx scores (n = 6 cells from 6 fish; Figure 3T).

In summary, *KalTA4u508* labels a specific group of pretectal neurons that are recruited during hunting initiation and which are capable of inducing naturalistic predatory behaviour in the absence of prey.

### Ablation of *KalTA4u508* pretectal neurons impairs hunting

To what extent are *KalTA4u508* pretectal neurons required for hunting? To address this question, we assessed hunting performance in freely swimming larvae provided with *Paramecia*, both before and after laser-ablation of *KalTA4u508* pretectal neurons (Figure 4A– C). To enable evaluation of the specificity of behavioural phenotypes, we also presented looming stimuli and drifting gratings to test visually evoked escape and optomotor response (OMR), respectively. Ablations were performed at 6 dpf in *KalTA4u508;UAS:mCherry* larvae and their efficacy was confirmed by reimaging the pretectum the following day. We estimated that ∼90% of the fluorescently labelled *KalTA4u508* pretectal population was typically ablated in both brain hemispheres (Figure 4D,E). Behaviour was tested both before (6 dpf) and after ablation (7 dpf) and control larvae underwent the same manipulations, other than laser-ablation, and were tested at the same time-points.

Analysis of prey consumption revealed that ablation of *KalTA4u508* pretectal neurons resulted in decreased hunting performance (Figure 4F). Further analysis revealed that this reduction in prey capture was associated with a reduced rate of hunting initiation (Figure 4G) as well as a reduction in the duration of hunting routines (Figure 4H). By contrast, we did not observe changes in average swim speeds, loom-evoked escapes or OMR performance (Figure 4I–L) and control larvae did not show changes in any of the tested behaviours between 6 and 7 dpf (Figure 4–supplement 2A–G). Together, these results indicate that *KalTA4u508* pretectal neurons are specifically required for normal initiation and maintenance of predatory behaviour.

Optogenetic stimulation of the anterior-ventral optic tectum (avOT) in *elavl3:itTA;Ptet:ChR2-YFP* transgenic larvae has been previously reported to induce convergent saccades and J-turns (Fajardo *et al.*, 2013). We examined ChR2-YFP expression in 6 dpf *elavl3:itTA;Ptet:ChR2-YFP* larvae and confirmed that the opsin is highly expressed in avOT as well as AF7 (Figure 4–supplement 1A,C), but is absent from *KalTA4u508* pretectal neurons (Figure 4–supplement 1E). The expression of ChR2 in AF7 raised the possibility that stimulation of the axon terminals of the prey-responsive RGCs that innervate this AF (Semmelhack *et al.*, 2014) might contribute to the hunting behaviour observed in this transgenic line. However, by crossing *elavl3:itTA;Ptet:ChR2-YFP* to the *lakritz* mutant, in which no RGCs are generated (Kay *et al.*, 2001), we found that retinally blind (*lak*^-/-^) transgenic animals were in fact more likely to display optogentically induced hunting than their sighted (*lak*^*+/+*^ or *lak*^+/-^) siblings (Figure 4–supplement 1,F–I and Video 3). In addition, responsive *lak*^-/-^ transgenics showed an increased probability of optogenetically induced hunting events, longer hunting routine durations and a substantial reduction in response latency (Figure 4– supplement 1J–L). These data are compatible with the conclusion of Fajardo *et al.* (2013), namely that optogenetic stimulation of avOT elicits hunting in *elavl3:itTA;Ptet:ChR2-YFP* and indicate that RGC stimulation (either visually with bright blue light, or optogenetically) interferes with induction of hunting responses.

To assess if AF7-pretectal circuits are required for such tectally induced hunting-like behaviour, we tested whether hunting could be evoked by optogenetic stimulation of avOT in larvae in which *KalTA4u508* pretectal cells were ablated. Following laser-ablation of *KalTA4u508* neurons in *elavl3:itTA;Ptet:ChR2-YFP;KalTA4u508;UAS:mCherry* larvae, we observed that the probability of optogenetically induced hunting was substantially reduced (Figure 4M). By contrast, control larvae showed no change in response probability between 6 and 7 dpf (Figure 4–supplement 2H).

In summary, these data indicate that *KalTA4u508* pretectal neurons contribute to the initiation and maintenance of natural hunting behaviour and are required for release of predatory behaviour by circuits in the anterior optic tectum.

## Discussion

In this study we combined multi-photon calcium imaging, single-cell optogenetic stimulation and laser-ablations to identify a population of pretectal neurons that commands hunting behaviour. Calcium imaging during naturalistic behaviour revealed that *KalTA4u508* pretectal neurons are recruited when larval zebrafish initiate hunting. Optogenetic activation of single *KalTA4u508* pretectal cells could release predatory behaviour in the absence of prey, and ablation of these cells impaired both natural hunting as well as hunting-like behaviour evoked by avOT stimulation. Based on functional and anatomical data, we propose that *KalTA4u508* pretectal cells comprise a command system linking visual perception of prey-like stimuli to activation of tegmental pattern generating circuits that coordinate specialised hunting motor programmes.

### A command system controlling predatory behaviour in AF7-pretectum

Our data support the idea that *KalTA4u508* pretectal neurons satisfy the criteria for a command system for induction of predatory behaviour. A command neuron has been defined as ‘a neuron that is both necessary and sufficient for the initiation of a given behaviour’ (Kupfermann and Weiss, 1978). Although a small number of striking examples of such cells have been identified in invertebrate models (Frost and Katz, 1996; Flood *et al.*, 2013), it is recognised that the ‘necessity’ criterion is unlikely to be fulfilled in larger nervous systems where individual neurons display functional redundancy (Yoshihara and Yoshihara, 2018). Thus, command systems (sometimes referred to as decision neurons, command-like neurons or higher-order neurons) have been defined as interneurons that are active in association with a well-defined behaviour and whose activity can induce that behaviour, but without the strict necessity requirement (Jing, 2009; Yoshihara and Yoshihara, 2018). *KalTA4u508* pretectal neurons satisfy these criteria. First, these cells are recruited during naturalistic hunting. By comparing neural activity in response versus non-response trials we were able to disambiguate ‘sensory’ activity, evoked by prey-like visual cues, from activity specifically associated with execution of hunting behaviour. We discovered that *KalTA4u508* pretectal neurons, located close to AF7, show minimal, if any, activity in response to prey-like stimuli but are reliably activated when larvae release convergent saccades at the commencement of hunting. Strikingly, optogenetic stimulation of single *KalTA4u508* neurons could evoke sustained hunting routines. Crucially, this induction occurred in the absence of any prey-like stimulus, indicating that these neurons directly command predatory behaviour and function downstream of the perceptual recognition of prey, rather than having a positive modulatory (‘motivating’) action on sensorimotor activity. This is one of the first examples of a behavioural action sequence induced by activation of a single neuron in a vertebrate.

Ablations targeting *KalTA4u508* pretectal neurons impaired, but did not eliminate, hunting. One contributing factor is likely to be that ablations were incomplete. We observed that 10% of labelled *KalTA4u508* neurons survived ablation and due to variegation of transgene expression (Akitake *et al.*, 2011), this is probably a lower bound on the size of the surviving pretectal population. Notably, the observation that hunting can be evoked by stimulation of only a single neuron suggests few surviving cells could in principle suffice to release predatory behaviour. Although our behavioural epistasis test revealed that hunting evoked by stimulation of anterior-ventral OT was strongly diminished in *KalTA4u508*-ablated animals, we do not rule out the possibility that there might be redundant, distributed circuitry involved in hunting initiation.

Overall, our data identify a discrete population of pretectal neurons that comprise a command system controlling predatory hunting.

### Neural circuitry controlling visually guided hunting

How might these pretectal neurons fit within a sensorimotor pathway controlling visually guided hunting (Figure 4N)? Current evidence suggests visual recognition of prey is mediated by tectal and/or pretectal circuits. The axon terminals of zebrafish ‘projection class 2’ retinal ganglion cells in AF7 appear tuned to prey-like stimuli (Semmelhack *et al.*, 2014) and a subpopulation of tectal neurons show non-linear mixed selectivity for conjunctions of visual features that best evoke predatory responses (Bianco and Engert, 2015). In accordance with these findings, we observed prey-responsive neurons in tectum and AF7-pretectum which were activated by small moving spots regardless of whether or not the animal produced a hunting response. Ablations of either AF7 or tectal neuropil have been shown to substantially impair hunting (Gahtan *et al.*, 2005; Semmelhack *et al.*, 2014), compatible with an important function for these retinorecipient regions in visual perception of prey.

Command systems are thought to sit at the sensorimotor ‘watershed’, linking sensory processing to activation of motor hierarchies. Our data support a circuit organisation where downstream of prey detection, the recruitment of *KalTA4u508* pretectal neurons might be the neural event that corresponds to the decision of the animal to initiate hunting. The dendritic arbours of these pretectal neurons lie in immediate apposition to RGC terminals in AF7, suggesting a biological interface for transforming visual sensory activity into premotor commands to release behaviour. Photoactivation data also suggests the AF7-pretectal region is closely interconnected with ipsilateral OT (Figure 2–supplement 1D), providing a further route for visual prey detectors in OT to recruit a hunting response. Consistent with this, our ablation data indicate that induction of hunting by stimulation of anterior-ventral OT (Fajardo *et al.*, 2013) requires *KalTA4u508* pretectal neurons.

Two morphological classes of *KalTA4u508* neuron were able to induce predatory behaviour with similar efficacy and motor kinematics, but opposite directional biases. Both morphologies appear similar to neurons previously identified in the AF7 region by Semmelhack *et al.* (2014). Contralaterally projecting neurons extended axon collaterals around the nMLF and oculomotor nuclei as well as in close apposition to reticulospinal neurons in the hindbrain. This projection pattern, as well as the identification of *KalTA4u508* neurons in adult brain sections, suggests these cells reside in the larval zebrafish APN. Their axonal projections provide a pathway by which pretectal commands could recruit tegmental motor pattern generators to produce the specialised eye and tail movements observed during hunting (McElligott and O’Malley D, 2005; Bianco *et al.*, 2011; Marques *et al.*, 2018). Optogenetic stimulation of ipsilaterally projecting *KalTA4u508* neurons evoked hunting routines in which the first orienting turn displayed an ipsilateral bias. It is possible that this arises from recruitment of tectal efferent pathways to the ipsilateral tegmentum, which have recently been suggested to mediate prey-directed orienting turns (Helmbrecht *et al.*, 2018).

Whilst it is remarkable that stimulation of single pretectal neurons could induce hunting-like behaviour in the absence of prey, natural hunting is a precise orienting behaviour directed towards a (visual) target. Optogenetically induced hunting routines involved swim bouts where turn angle fell within the lower portion of the distribution measured during prey hunting. A probable explanation is that in the presence of prey, appropriate steering signals derive from OT to guide lateralised orienting turns. Compact tectal assemblies show premotor activity immediately preceding hunting initiation and are anatomically localised in relation to the directionality of hunting responses (Bianco and Engert, 2015). Furthermore, focal stimulation of the retinotopic tectal map has long been known to evoke goal-directed orienting responses including predatory manoeuvres (Herrero *et al.*, 1998; Bels *et al.*, 2012). We hypothesise that pretectal activity releases predatory behaviour and operates in parallel with prey-directed steering signals, most likely from OT.

Command systems have been identified that evoke both discrete actions (Korn and Faber, 2005) as well as entire behavioural programmes (Flood *et al.*, 2013). Constant stimulation of pretectal *KalTA4u508* neurons produces extended hunting sequences that commenced with saccadic eye convergence followed by serial execution of multiple swim bouts during which elevated ocular vergence – a hallmark of predatory behaviour – was maintained. This suggests that the function of pretectal activity is to command the overall hunting programme and that individual component actions (*i.e.* discrete tracking swim bouts) might be coordinated by downstream motor pattern generating circuits.

### Control and modulation of predatory behaviour

In recent years, several studies in rodents have demonstrated that pathways converging on the periaqueductal grey (PAG) can potently modulate predatory behaviour. The central amygdala (CeA) displays activity changes during hunting (Comoli *et al.*, 2005) and optogenetic activation of the CeA→ventral PAG pathway motivates prey pursuit (Han *et al.*, 2017). Stimulation of the medial preoptic area to vPAG pathway promotes acquisition/handling (grabbing, biting) of objects, including prey (Park *et al.*, 2018), and activation of a GABAergic projection from lateral hypothalamus to lateral/ventrolateral PAG motivates attack on prey (Li *et al.*, 2018). In these studies, predatory behaviour was induced in the presence of prey/prey-like stimuli, suggesting these pathways serve to motivate, rather than command, hunting. In future studies, it will be interesting to examine the roles of mammalian thalamic pretectal nuclei as well as the superior colliculus, in commanding and directing predatory responses. Command systems represent a key circuit node for integration of sensory information with internal state signals (Flood *et al.*, 2013). In the context of vertebrate hunting, it will be important to elucidate the circuit mechanisms by which regions such as lateral hypothalamus modulate the recruitment probability of hunting command systems, enabling animals to vary the expression of predatory behaviour in accordance with internal drives, experience and competing behavioural demands.

## Acknowledgements

The authors thank members of the Bianco lab, David Attwell, Tiago Branco, Tara Keck and Steve Wilson for helpful discussions and critical feedback on the manuscript and UCL Fish Facility staff for fish care and husbandry. We thank Richard Poole for help with the laser-ablation system and Claire Wyart for sharing the *Tg(UAS:GCaMP6f,cryaa:mCherry)*^*icm06Tg*^ line prior to publication. Joanna Lau kindly provided the schematic in Figure 1A and Pedro Henriques provided the anatomical mask of the *chata*^*+*^ nucleus isthmi in ZBB coordinates. P.A. was supported by a Sir Henry Wellcome Postdoctoral Fellowship (204708/Z/16/Z). I.H.B. was supported by a Sir Henry Dale Fellowship from the Royal Society & Wellcome Trust (101195/Z/13/Z) and a UCL Excellence Fellowship.

## Author Contributions

Conceptualization and Experimental Design: P.A. and I.H.B.; Data Collection: P.A. with exception of adult neuroanatomy data collected by M.F.; Analysis: P.A.; Writing: P.A. and I.H.B.; Funding Acquisition: P.A and I.H.B.

## Materials and Methods

### Experimental model and transgenic lines

Animals were reared on a 14/10 h light/dark cycle at 28.5°C. For all experiments, we used zebrafish larvae homozygous for the *mitfa*^*w2*^ skin-pigmentation mutation (Lister *et al.*, 1999). Larvae used for pan-neuronal Ca^2+^ imaging experiments were double-transgenic *Tg(elavl3:H2B-GCaMP6s)*^*jf5Tg*^ (Vladimirov *et al.*, 2014) and *Tg(atoh7:gapRFP)*^*cu2Tg*^ (Zolessi *et al.*, 2006). For AF7-pretectal Ca^2+^ imaging, larvae were double-transgenic for *Tg(– 2.5pvalb6:KalTA4)*^*u508Tg*^ [*i.e. Tg(KalTA4u508*); generated in this study, see below] and either *Tg(UAS:GCaMP6f,cryaa:mCherry)*^*icm06Tg*^ (Knafo *et al.*, 2017) or *Tg(UAS:jGCaMP7f)*^*u341Tg*^ (generated in this study, see below). Larvae used to determine whether *KalTA4u508* neurons reside in AF7-pretectum were triple-transgenic *Tg(KalTA4u508), Tg(UAS-E1b:NfsB-mCherry)*^*jh17Tg*^ (Davison *et al.*, 2007) and *TgBAC(slc17a6b:loxP-DsRed-loxP-GFP)*^*nns14Tg*^ (Koyama *et al.*, 2011). Larvae used for AF7 dendritic stratification analyses were triple-transgenic *Tg(KalTA4u508), Tg(UAS:RFP)*^*tpl2Tg*^ (Auer *et al.*, 2014), and *Tg(atoh7:GFP)*^*rw021Tg*^ (Masai *et al.*, 2003). Fish used for mapping of cell location in the adult pretectum were triple-transgenic *Tg(KalTA4u508), Tg(UAS:GCaMP6f,cryaa:mCherry)*^*icm06Tg*^ and *Tg(atoh7:gapRFP)*^*cu2Tg*^. Larvae used for photo-activatable GFP labelling were *Tg(Cau.Tuba1:c3paGFP)*^*a7437Tg*^ (Bianco *et al.*, 2012). Larvae used for single cell labelling and optogenetic stimulation of AF7-pretectal cells were double-transgenic *Tg(KalTA4u508)* and *Tg(elavl3:H2B-GCaMP6s)*^*jf5Tg*^. Larvae used for pretectal cell ablations and free-swimming behaviour analyses were triple-transgenic *Tg(KalTA4u508), Tg(UAS-E1b:NfsB-mCherry)*^*jh17Tg*^ (Davison *et al.*, 2007) and *Tg(elavl3:ITETA-PTET:Cr.Cop4-YFP)*^*fmi2Tg*^ (Fajardo *et al.*, 2013). Larvae used for optogenetic stimulation of the avOT were double transgenic *Tg(atoh7:gapRFP)*^*cu2Tg*^ and *Tg(elavl3:ITETA-PTET:Cr.Cop4-YFP)*^*fmi2Tg*^ with either homozygous, heterozygous or no mutation of the *atoh7*^*th241*^ gene (Kay *et al.*, 2001). All larvae were fed *Paramecia* from 4 dpf onward. Animal handling and experimental procedures were approved by the UCL Animal Welfare Ethical Review Body and the UK Home Office under the Animal (Scientific Procedures) Act 1986.

### 2-photon calcium imaging and behavioural tracking

The procedure was very similar to that described in Bianco and Engert (2015). Larval zebrafish were mounted in 3% low-melting point agarose (Sigma-Aldrich) at 5 dpf or 6 dpf and allowed to recover overnight before functional imaging at 6 dpf or 7 dpf. Imaging was performed using a custom-built 2-photon microscope [Olympus XLUMPLFLN 20× 1.0 NA objective, 580 nm PMT dichroic, bandpass filters: 510/84 (green), 641/75 (red) (Semrock), Coherent Chameleon II ultrafast laser]. Imaging was performed at 920 nm with average laser power at sample of 5–10 mW. For imaging of *Tg(elavl3:H2B-GCaMP6s)* larvae, images (500×500 pixels, 0.61 *µ*m/px) were acquired by frame scanning at 3.6 Hz and for each larva 10–14 focal planes were acquired with a z-spacing of 8 *µ*m. For imaging of *Tg(KalTA4u508;UAS:GCaMP6f)* or *Tg(KalTA4u508;UAS:jGCaMP7f)* larvae, the same image size and scanning rate were used but 5–6 focal planes with a z-spacing of 5 *µ*m were acquired for each larva. Visual stimuli were back-projected (Optoma ML750ST) onto a curved screen placed in front of the animal at a viewing distance of ∼7 mm while a second projector provided constant background illumination below the fish. A coloured Wratten filter (Kodak, no. 29) was placed in front of both projectors to block green light from the PMT. Visual stimuli were designed in MATLAB using Psychophysics toolbox (Brainard, 1997). For all experiments, stimuli were presented in a pseudo-random sequence with 30 s inter-stimulus interval. Stimuli comprised 5° or 16°, dark or bright spots moving at 30°/s either left→right or right→left across ∼200° of frontal visual space. Bright/dark spots had Weber contrast of 1/-1, respectively.

In addition, 3 s whole-field bright/dark flashes were presented. Eye movements were tracked at 60 Hz under 720 nm illumination using a FL3-U3-13Y3M-C camera (Point Grey) that imaged through the microscope objective. Tail movements were imaged at 430 Hz under 850 nm illumination using a sub-stage GS3-U3-41C6NIR-C camera (Point Grey). Microscope control, stimulus presentation and behaviour tracking were implemented using custom LabVIEW and MATLAB software.

### Calcium imaging analysis

All calcium imaging data analysis was performed using custom-written MATLAB scripts. Motion correction of fluorescence imaging data was performed as per Bianco and Engert (2015). Regions of interest (ROIs) corresponding to cell nuclei were extracted using the cell detection code from Kawashima *et al.* (2016). The time-varying fluorescence signal *F(t)* for each cell was extracted by computing the mean value of all pixels within the corresponding ROI binary mask at each time-point (imaging frame). The proportional change in fluorescence (*ΔF/F*_*0*_) at time *t* was calculated as

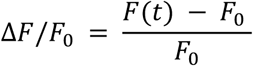

where *F*_*0*_ is a reference fluorescence value, taken as the median of *F(t)* during the 30 frames prior to all visual stimulus presentations.

We used multilinear regression to model the fluorescent timeseries of each imaged neuron (ROI) in terms of simultaneously recorded kinematic predictors (‘regressors’). Regressors were generated for oculomotor and locomotor variables (7 eye and 3 tail kinematics, see Supplementary File 1) by convolving time-series vectors for the relevant kinematic with a calcium impulse response function [CIRF, approximated as the sum of a fast-rising exponential, tau 20 ms, and a slow-decaying exponential, tau 420 ms for GCaMP6f and jGCaMP7f or 3 s for H2B-GCaMP6s; (Miri *et al.*, 2011)]. To account for delays between neural activity and behaviour and/or indicator dynamics, we time-shifted the regressor matrix relative to the fluorescent response variable so as to minimise the residual squared error of an ordinary least squares regression model. We used elastic-net regularised regression to improve interpretability and prediction accuracy in the presence of multicollinearity between the regressors (Zou and Hastie, 2005). Elastic net models were fit using the ‘glmnet’ package for MATLAB (Qian *et al.*, 2013) and hyperparameters controlling the ratio of L1 vs. L2 penalty (alpha) and the degree of regularization (lambda) were selected to minimise ten-fold cross-validated squared error. Model coefficients were then used to construct visuomotor vectors.

Visuomotor vectors (VMVs) were generated for each neuron by concatenating (a) the integral of *ΔF/F*_*0*_ in response to each visual stimulus (12 s time window from stimulus onset, mean integral across stimulus presentations) for presentations in which no eye convergence was performed by the larva (components 1–10); (b) multilinear regression coefficients (*β* values) for eye, tail and motion correction regressors (components 11–21). VMVs from all imaged neurons were assembled into a matrix and each component was normalised across cells by dividing each column by its standard deviation.

VMV clustering was performed using a two-step procedure. First, we performed hierarchical agglomerative clustering of VMVs using a correlation distance metric (Bianco and Engert, 2015). For this first step, we selected only neurons that either exhibited strong visual responses (specifically, the maximum value of components 1–10 had to be within the top 5^th^ percentile of such values across all neurons) or was well modelled in terms of behavioural kinematics (R^2^ had to be within the top 5^th^ percentile of cross-validated R^2^ across all neurons). The centroids of clusters generated in this step (correlation threshold, 0.7) constituted a set of archetypal response profiles. Next, the VMVs of the remaining neurons were assigned to the cluster with the closest centroid (within a correlation distance threshold of 0.7).

To assign cluster identities to *KalTA4u508* pretectal neurons (*e.g.* Figure 2J and 3T), VMVs were generated as described above and the same assignment strategy and correlation distance threshold (0.7) were used. Note that normalisation of components was performed using the standard deviations computed for the initial matrix of VMVs.

Hunting Index (HIx) scores were calculated for each cell as follows. For each hunting response, convergence-triggered activity was measured by computing the mean of z-scored GCaMP fluorescence in a time window (±1 s) centred on the convergent saccade, *x*_*Ri*_. Next, activity was measured at the same time during non-response trials in which the same visual stimulus was presented. The difference between *x*_*Ri*_ and the mean of non-response activity, *μ*_*NR*_, was computed:

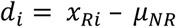

HIx scores were computed as the mean of these *d*_*i*_ distance values across all response trials during which the cell was imaged. CMI values were computed separately for convergence events paired with leftwards tail movements, rightwards tail movements, and symmetrical/no tail movements.

### 3D image registration

Registration of image volumes was performed using the ANTs toolbox version 2.1.0 (Avants *et al.*, 2011). Images were converted to the NRRD file format required by ANTs using ImageJ. As an example, to register the 3D image volume in ‘fish1_01.nrrd’ to the reference brain ‘ref.nrrd’, the following parameters were used:

~~~
antsRegistration -d 3 -float 1 -o [fish1_, fish1_Warped.nii.gz]
-n BSpline -r [ref.nrrd, fish1_01.nrrd, 1] -t Rigid[0.1] -m
GC[ref.nrrd, fish1_01.nrrd, 1, 32, Regular, 0.25] -c
[200×200×200×0,1e-8, 10] -f 12×8×4×2 -s 4×3×2×1 -t Affine[0.1] -
m GC[ref.nrrd, fish1_01.nrrd, 1, 32, Regular, 0.25] -c
[200×200×200×0,1e-8, 10] -f 12×8×4×2 -s 4×3×2×1 -t SyN[0.1, 6, 0]
-m CC[ref.nrrd, fish1_01.nrrd, 1, 2] -c [200×200×200×200×10,1e-
7, 10] -f 12×8×4×2×1 -s 4×3×2×1×0
~~~

The deformation matrices computed above were then applied to any other image channel N of fish1 using:

~~~
antsApplyTransforms -d 3 -v 0 -float -n BSpline -i fish1_01.nrrd
-r ref.nrrd -o fish1_0N_Warped.nii.gz -t fish1_1Warp.nii.gz -t
fish1_0GenericAffine.mat
~~~

All brains were registered onto the ZBB brain atlas (1×1×1 xyz *µ*m/px) (Marquart *et al.*, 2015; Marquart *et al.*, 2017) and onto a high-resolution *Tg(elavl3:H2B-GCaMP6s)* reference brain (0.76×0.76×1 xyz *µ*m/px, mean of 3 larvae), with some differences between experiments:

- For functional calcium imaging volumes, a three-step registration was used: the imaging volume, composed of 10–14 image planes (500×500 px, 0.61 *µ*m/px, 8 *µ*m z-spacing), was first registered to a larger volume of the same brain acquired at the end of the experiment (1 *µ*m z-spacing), using affine and warp transformations. Then, the larger volume was registered to the Hi-Res *Tg(elavl3:H2B-GCaMP6s)* reference brain. Because the high-resolution volume had already been registered onto the ZBB atlas, the transformations were concatenated to bring the functional imaging volume to the ZBB atlas (calcium imaging stack → post-imaging stack → Hi-Res → ZBB).
- The brain regions displayed in Figure 2F,H, and Figure 2–supplement 1A,B correspond to volumetric binary image masks in the ZBB atlas that have been registered to the Hi-Res *Tg(elavl3:H2B-GCaMP6s)* reference brain using the ZBB *Tg(elavl3:H2B-RFP)* volume and performing affine and warp transformations (ZBB *elavl3:H2B-RFP* → Hi-Res).
- For the registration displayed in Figure 2A,B of *KalTA4u508* neurons to the Hi-Res *Tg(elavl3:H2B-GCaMP6s)* reference brain, the imaging volume was registered to the ZBB *Tg(vglut:DsRed)* volume [previously registered to the Hi-Res reference brain (ZBB *elavl3:H2B-RFP* → Hi-Res] using the vglut2a channel [*TgBAC(slc17a6b:loxP-DsRed-loxP-GFP)*] acquired in parallel with *Tg(KalTA4u508*;*UAS-E1b:NfsB-mCherry)* imaging and performing affine and warp transformations.
- For single *KalTA4u508* neuron tracing experiments, the imaging volume was first registered to the Hi-Res *Tg(elavl3:H2B-GCaMP6s)* reference brain using the H2B-GCaMP6s channel acquired in parallel with *Tg(KalTA4u508);UAS-CoChR-tdTomato-injected* imaging and performing affine and warp transformations. Transformations were then concatenated to bring the imaging volume and associated neuron tracing (see below) to the ZBB atlas (imaging stack → Hi-Res → ZBB).
- Imaging volumes related to photo-activation of PA-GFP were registered to a whole-brain reference from a 6 dpf *Tg(α-tubulin:C3PA-GFP)* larva in which no photo-activation was performed, using affine and warp transformations. The *Tg(α-tubulin:C3PA-GFP)* reference volume was then registered to the ZBB atlas. The photo-activation volume was transported to the Hi-Res *Tg(elavl3:H2B-GCaMP6s)* reference by concatenating the transformations (photo-activation stack → *Tg(α-tubulin:C3PA-GFP)* reference → ZBB → Hi-Res).

All registration steps were manually assessed for global and local alignment accuracy. All brain regions referred to in this paper correspond to the volumetric binary image masks in the ZBB atlas, with the exception of regions in the anterior-ventral optic tectum, AF7-pretectum, and cholinergic nucleus isthmi. These image masks, in ZBB reference space, can be downloaded as Supplementary File 2–4.

### DNA cloning and transgenesis

To generate the *UAS:CoChR-tdTomato* DNA construct used for single cell labelling and optogenetic stimulations, the coding sequence of the blue light-sensitive opsin CoChR (from *pAAV-Syn-CoChR-GFP*) and the red fluorescent protein tdTomato (from *pAAV-Syn-Chronos-tdTomato*) were cloned in frame into a UAS Tol1 backbone (*pT1UciMP*). The *pAAV-Syn-CoChR-GFP* and *pAAV-Syn-Chronos-tdTomato* plasmids were gifts from Edward Boyden (Addgene plasmid # 59070 and # 62726, respectively) (Klapoetke *et al.*, 2014). The *pT1UciMP* plasmid was a gift from Harold Burgess (Addgene plasmid # 62215) (Horstick *et al.*, 2015). The cloning was achieved using the In-Fusion HD Cloning Plus CE kit (Clontech) with the following primers:

- CoChR_fw, CTCAGCGTAAAGCCACCATGCTGGGAAACG
- CoChR_rev, TACTACCGGTGCCGCCACTGT
- CoChR_tdT_fw, ACAGTGGCGGCACCGGTAGTA
- tdT_rev, CTAGTCTCGAGATCTCCATGTTTACTTATACAGCTCATCCATGCC

To generate the *UAS:jGCaMP7f* DNA construct used for creating the *Tg(UAS:jGCaMP7f)*^*u341Tg*^ line, the coding sequence of the genetically encoded calcium indicator jGCaMP7f (from *pGP-CMV-jGCaMP7f*) was cloned into the *pT1UciMP* UAS Tol1 backbone. The *pGP-CMV-jGCaMP7f* plasmid was a gift from Douglas Kim (Addgene plasmid # 104483) (Dana *et al.*, 2018). As above, the cloning was achieved using the In-Fusion HD Cloning Plus CE kit (Clontech) with the following primers:

- UAS_jGCaMP7_fw, CGTAAAGCCACCATGGGTTCTCATC
- UAS_jGCaMP7_rev, CTCGAGATCTCCATGTTTACTTCGCTGTCATCATTTGTACAAAC

To generate the *Tg(UAS:jGCaMP7f)* line, purified *UAS:jGCaMP7f* DNA constructs (35 ng/*µ*l) were co-injected with Tol1 transposase mRNA (80 ng/*µ*l) into *Tg(KalTA4u508)* zebrafish embryos at the early one-cell stage. Transient expression, visible as jGCaMP7f fluorescence, was used to select injected embryos that were then raised to adulthood. *Tol1* transposase mRNA was prepared by *in vitro* transcription from NotI-linearised *pCS2-Tol1.zf1* plasmid using the SP6 transcription mMessage mMachine kit (Life Technologies). The *pCS2-Tol1.zf1* was a gift from Harold Burgess (Addgene plasmid # 61388) (Horstick *et al.*, 2015). RNA was purified using the RNeasy MinElute Cleanup kit (Qiagen). Germ line transmission was identified by mating sexually mature adult fish to *mitfa* fish and, subsequently, examining their progeny for jGCaMP7f fluorescence. Positive embryos from a single fish were then raised to adulthood. Once this second generation of fish reached adulthood, positive embryos from a single ‘founder’ fish were again selected and raised to adulthood to establish a stable *Tg(KalTA4u508;UAS:jGCaMP7f)* double transgenic line.

The *Tg(–2.5pvalb6:KalTA4)*^*u508Tg*^ [*i.e. Tg(KalTA4u508*)] line was isolated as follows. First, we used Gateway cloning (Invitrogen) to construct an expression vector in which ∼2.5 kb of zebrafish genomic sequence upstream of the *pvalb6* gene start codon was placed upstream of the KalTA4 (Distel *et al.*, 2009) open reading frame. The genomic sequence was cloned using the following primers and Phusion PCR polymerase (Thermo Fisher Scientific):

- fwd: GGGGACAAGTTTGTACAAAAAAGCAGGCTggatggtgggccaaatcaaaggctac
- rev: GGGGACCACTTTGTACAAGAAAGCTGGGTggaacgagaccggcaacacacag

(where capital letters indicate the attB1/B2 extension sequences).

The expression vector was then micro-injected into one-cell stage *Tg(UAS-E1b:Kaede)*^*s1999t*^ (Davison *et al.*, 2007) embryos at 30 ng/*µ*l along with *tol2* mRNA (30 ng/*µ*l) and adult fish were screened for germline transmission by outcrossing as described above. This expression vector generated a wide range of expression patterns, one of which labelled AF7-pretectal neurons as reported here (allele *u508Tg*).

### Immunohistochemistry

#### Larvae

Samples were fixed overnight in 4% paraformaldehyde (PFA) in 0.1 M phosphate buffered saline (PBS, Sigma-Aldrich) and 4% sucrose (Sigma-Aldrich) at 4°C. Brains were manually dissected with forceps prior to immunostaning. First, dissected brains were permeabilised by incubation in proteinase-K (40 *µ*g/ml) in PBS with 1% Triton-X100 (PBT, Sigma-Aldrich) for 15 minutes. This was followed by 3× 5 min washes in PBT, 20 min fixation in 4% PFA at room temperature and 3× 5 min washes in PBT. Second, brains were incubated in block solution (2% goat serum, 1% DMSO, 1% BSA in PBT, Sigma-Aldrich) for 2 h. Subsequently, brains were incubated in block solution containing primary antibodies overnight, followed by 6× 1 h washes in PBT on a slowly rotating shaker. Third, brains were incubated in block solution containing secondary antibodies overnight, followed by 6× 1 h washes in PBT. Finally, PBT was rinsed out by doing washes in PBS and brains were stored at 4°C. Imaging was performed using the two-photon microscope described above at 790 nm. Primary antibodies were: rabbit anti-GFP (AMS Biotechnology, TP401, dilution 1:1000) and mouse anti-ERK (Cell Signaling Technology, 9102, p44/42 MAPK (Erk1/2), dilution 1:500). Secondary antibodies were: goat anti-rabbit Alexa Fluor 488-conjugated (Thermo Fisher Scientific, A-11034, dilution 1:200), and goat anti-mouse Alexa Fluor 594-conjugated (Thermo Fisher Scientific, A-11005, dilution 1:200).

#### Adults

*Tg(KalTA4u508;UAS:GCaMP6f;atoh7:gapRFP)* fish (3 months old) were deeply anesthetized in 0.2% tricaine (MS222, Sigma) and fixed in 4% paraformaldehyde (PFA) for 24 h at 4°C. Brains were then carefully dissected under a stereomicroscope and transferred to saline phosphate buffer (PBS), where they were maintained for at least half an hour. Two different procedures were used for sectioning the brains. For cryostat sectioning, brains were cryopreserved, embedded in Tissue Tek OCT compound (Sakura Finetek) and frozen using liquid nitrogen cooled methylbutane. Transverse sections of the brain (12 *µ*m thick) were obtained using a cryostat and collected in gelatine-coated slides. For vibratome sectioning, brains were first embedded into 3% agarose in PBS. Transverse sections of the brains (100 *µ*m thick) were obtained using a vibratome and transferred to PBS in microtubes. Immunostaining was performed by either adding solutions onto the cryostat sections or changing the solutions inside the microtubes. Sections were incubated first in normal goat serum (Sigma, dilution 1:10) in PBS with 0.5% Triton for 1 h at room temperature, and then with a cocktail of two primary antibodies (rat anti-GFP, Nacalai Tesque, 04404-26, dilution 1:1000, and rabbit anti-RFP, MBL International, PM005, dilution 1:1000) for 24 h at room temperature. Next, after three washes in PBS, brain sections were incubated with a cocktail of two secondary antibodies (goat anti-rat Alexa 488, Thermo Fisher Scientific, A-11006, dilution 1:500, and goat anti-rabbit Alexa 568, Thermo Fisher Scientific, A-11011, dilution 1:500) for 1 h at room temperature. Sections were washed in PBS, mounted with 50% glycerol in PBS and imaged using a Nikon A1R confocal microscope equipped with a Nikon Plan Fluor 20× 0.50 NA objective. Excitation light was provided by an argon ion multichannel laser tuned to 488 nm (green channel), and a 561 nm diode laser (red channel).

### Single cell labelling

To label individual *KalTA4u508* neurons, *UAS:CoChR-tdTomato* DNA constructs were injected into 1–4 cell stage *Tg(KalTA4u508)* embryos. Plasmid DNA was purified using midi-prep kits (Qiagen) and injected at a concentration of 30 ng/*µ*l in distilled water. Larvae (4 dpf) were then screened for CoChR-tdTomato expression. Only larvae showing expression in a single *KalTA4u508* pretectal neuron were subsequently used for optogenetic stimulations and neuronal tracing experiments. Single cell morphologies were traced using the Simple Neurite Tracer plugin for ImageJ (Longair *et al.*, 2011).

### Anatomical analyses

Cell density of neurons belonging to hunting-initiation clusters (25–28) was computed in the following way. First, we obtained the soma 3D coordinates of all cluster 25–28 neurons following anatomical registration to the high-resolution *Tg(elavl3:H2B-GCaMP6s)* reference brain. Then, we computed the local cell density at each soma location using an adaptive Gaussian-based kernel density estimate (Breiman *et al.*, 1977), with the bandwidth at each point constrained to be proportional to the *k*th nearest neighbour distance where:

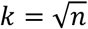

and *n* is the number of neurons (n = 6,630 cells from 8 fish). To compute the kernel density estimate, we used a MATLAB-based toolbox developed by Alexander Ihler (www.ics.uci.edu/~ihler/code/kde.html). Images in Figure 2A–C represent volume projections in which hunting-initiation neurons are colour-coded according to local cell density.

Neurite stratification and axon projection profiles in Figure 2D,G were obtained by measuring fluorescence intensity values along the axes indicated on figure panels using ImageJ ‘Line’ and ‘Plot Profile’ tools. For each image channel, a maximum intensity projection image was generated before measuring fluorescence intensity. Each intensity profile *i* was then rescaled to generate a profile, *I*, ranging from 0 to 1 as follows:

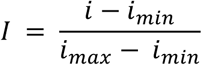

where *i*_*min*_ and *i*_*max*_ are the minimum and maximum values of profile *i*, respectively.

### Photo-activation of PA-GFP

Larvae (5 dpf) homozygous for the *Tg(α-tubulin:C3PA-GFP)* transgene were anaesthetised and mounted in 2% low-melting temperature agarose. The same custom-built 2-photon microscope used for functional imaging was used to photo-activate PA-GFP in a small region (9×9 *µ*m) containing cell bodies located in AF7-pretectum. The photo-activation site was selected by imaging the brain at 920 nm. Photo-activation was performed by continuously scanning at 790 nm (5 mW at sample) for 4 min. Larvae were then unmounted and allowed to recover. At 7 dpf, an image stack (1200×800 px, 0.38 *µ*m/px, ∼200 *µ*m z-extent) was acquired at 920 nm covering a large portion of the midbrain, tegmentum and hindbrain. Axonal projections were traced using the Simple Neurite Tracer plugin for ImageJ (Longair *et al.*, 2011).

### Monitoring of free-swimming behaviour

The same behavioural tracking system was used for both optogenetic stimulations and assessment of visuomotor behaviours with some differences. Images were acquired under 850 nm illumination using a high-speed camera [Mikrotron, EoSens CL MC1362, 250 Hz (*optogenetic stimulations*) or 700 Hz (*visuomotor behaviours assessment*), 500 *µ*s shutter-time) equipped with a machine vision lens (Fujinon HF35SA-1) and an 850 nm bandpass filter to block visible light. In all experiments, larvae were placed in the arena and allowed to acclimate for around 2 min before starting experiments.

#### Optogenetic stimulations

Larvae were placed in a petri dish with a custom-made agarose well (28 mm diameter, 3 mm depth) filled with fish facility water. Blue light was delivered across the whole arena from above using a 470 nm LED (OSRAM Golden Dragon Plus, LB W5AM). ‘LED-On’ trials included 7 s periods of continuous blue light illumination (0.443 ± 0.001 mW/mm^2^, mean ± SD), interleaved with ‘no-stimulation’ trials in which no blue light was provided. Both ‘LED-On’ and ‘no-stimulation’ trials lasted 8 s. A minimum of 10 ‘LED-On’ trials were acquired for each fish. LED stimulus presentation and camera control were implemented using custom software written in LabVIEW (National Instruments).

#### Assessment of visuomotor behaviours

Larvae were placed in a 35 mm petri dish filled with 3.5 ml of fish facility water. Visual stimuli were projected onto the arena from below using an AAXA P2 Jr Pico Projector via a cold mirror. Visual stimuli were designed using Psychophysics Toolbox (Brainard, 1997). Looming stimuli expanded from 10–100° with L/V ratio of 255 ms (Dunn *et al.*, 2016). Optomotor gratings had a period of ∼10 mm and moved at 1 cycle/s. Optomotor gratings and looming spots were presented in egocentric coordinates such that directional gratings always moved 90° to left or right sides with respect to fish orientation and looming spots were centred 5 mm away from the body centroid and at 90° to left or right. Stimuli were presented in pseudo-random order with an inter-stimulus interval of minimum 60 s. Stimuli were only presented if the body centroid was within a predefined central region (11 mm from the edge of the arena). If this was not the case, a concentric grating was presented that moved towards the centre of the arena to attract the fish to the central region. At the beginning of each experiment, 60 *Paramecia* were added to arena. Each experiment typically lasted ∼1 hour. Final *Paramecia* numbers were counted manually from full-frame video data from the final 10 s of each experiment and adjusted for consumption in 60 min [multiplying by 60/experiment duration (min)]. During experiments, eye and tail kinematics were tracked online as described below. Camera control, online tracking and stimulus presentation were implemented using custom software written in LabVIEW (National Instruments) and MATLAB (MathWorks).

### Analyses of free-swimming behaviour

Data analysis was performed using custom software written in LabVIEW (National Instruments) and MATLAB (MathWorks). Eye and tail kinematics were tracked offline for optogenetic experiments, and online for assessment of visuomotor behaviours with some differences. First, images were background-subtracted using a background model generated over 8 s in which the larva was moving (*offline tracking*), or a continuously updated background model (*online tracking*). Next, images were thresholded and the body centroid was found by running a particle detection routine for binary objects within suitable area limits. For online tacking, eye centroids were detected using a second threshold and particle detection procedure with the requirement that these centroids were in close proximity to the body centroid. For offline tacking, eye centroids were detected using a particle detection procedure that uses both binary and greyscale images to identify the two centroids within suitable area limits that had the lowest mean intensity values. For online tracking, body and eye orientations were computed using second- and third-order image moments. For offline tracking, body orientation was computed as the angle of the vector formed by the centre of mass of the body centroid (origin) and the midpoint between the eye centroids. Eye orientation was computed as the angle between the major axis of the eye and the body orientation vector. Vergence angle was computed as the difference between the left and right eye angles. The tail was tracked by performing consecutive annular line-scans, starting from the body centroid and progressing towards the tip of the tail so as to define 9 equidistant x-y coordinates along the tail. Inter-segment angles were computed between the 8 resulting segments. Reported tail curvature was computed as the sum of these inter-segment angles. Rightward bending of the tail is represented by positive angles and leftward bending by negative angles. To identify periods of high ocular vergence, which represent hunting routines, a vergence angle threshold was computed for each fish by fitting a two-term Gaussian model to its vergence angle distribution. A fish was considered to be hunting if vergence angle exceeded this vergence threshold. For experiments assessing visuomotor behaviours, the vergence angle distribution was invariably bimodal and the vergence threshold was computed as one standard deviation below the centre of the higher angle Gaussian. For optogenetic experiments, in cases where the vergence angle distribution was not bimodal, a fixed vergence threshold of 55° was used.

#### Optogenetic experiments

Response probability was computed as the fraction of LED-On trials in which at least one hunting routine (*i.e.* period with ocular vergence above threshold) was detected during the 7 s stimulation period. Similarly, for no-stimulation trials, response probability is the fraction of trials in which at least one hunting-like routine was detected. Response latency for LED-On trials was calculated from light stimulus onset. Swim bouts were identified using velocity thresholds (800°/s for bout onset, 200°/s for bout offset) applied to smoothed absolute tail angular velocity traces. Tail beat frequency was computed as the reciprocal of the mean full-cycle period during a swim bout. Tail vigour is computed by integrating absolute tail angular velocity (smoothed with a 40 ms box-car filter) over the first 120 ms of a swim bout. Bout asymmetry measures the degree to which tail curvature during a bout shows the same laterality as that determined during the first half-beat. It is computed as the fraction of time points in which the sign of tail angle matches the direction of the first half beat. This metric is high for hunting related J-turns but close to zero for forward swims. For each bout, the fraction of total curvature localised to the distal third of the tail was computed for the first half beat.

#### Assessment of visuomotor behaviours

Escape responses to loom stimuli were identified if the instantaneous speed of the body centroid exceeded 75 mm/s. An optomotor response gradient [OMR turn rate (°/s)] was calculated for each presentation as the total change in orientation during the stimulus presentation divided by the duration of the presentation [for leftwards OMR stimuli, the OMR turn rate (typically negative) was multiplied by –1 to group the data with rightwards OMR stimuli]. Mean swim speed was calculated as the total distance covered by the larva in the central region of the arena divided by the total time spent in this region.

### Laser ablations

*KalTA4u508* pretectal neurons were targeted for ablation in 6 dpf *Tg(KalTA4u508;UAS:mCherry;elavl3:itTA;Ptet:ChR2-YFP)* larvae, which were anaesthetized using MS222 and mounted in 1% low-melting temperature agarose (Sigma-Aldrich). Ablations were performed using a MicroPoint system (Andor) attached to a Zeiss Axioplan-2 microscope equipped with a Zeiss Achroplan water-immersion 63× 0.95 NA objective. A pulsed nitrogen-pumped tunable dye laser (Coumarin-440 dye cell) was focused onto individual *KalTA4u508* neurons and pulses were delivered at a frequency of 10 Hz for 60– 120 s. All visible *KalTA4u508* neurons in both hemispheres were targeted for ablation and cell damage was confirmed under DIC optics. Larvae were then unmounted and allowed to recover overnight. Control larvae were mounted in 1% low-melting temperature agarose and underwent the same manipulations as ablated larvae except for laser-ablation. Pre- and post-ablation image stacks were acquired with a 2-photon microscope at 800 nm (800×800 px, 0.38 *µ*m/px, ∼40 *µ*m z-extent). Cell counting before and after ablations was performed manually in ImageJ using the multi-point tool.

### Quantification and Statistical Analysis

Statistical analyses were performed in Prism 8 (GraphPad) and MATLAB R2017b (MathWorks). Statistical tests, p-values, N-values, and additional information are reported in Supplementary File 1. All tests were two-tailed and were chosen after data were tested for normality and homoscedasticity.

**Figure 1–supplement 1.**
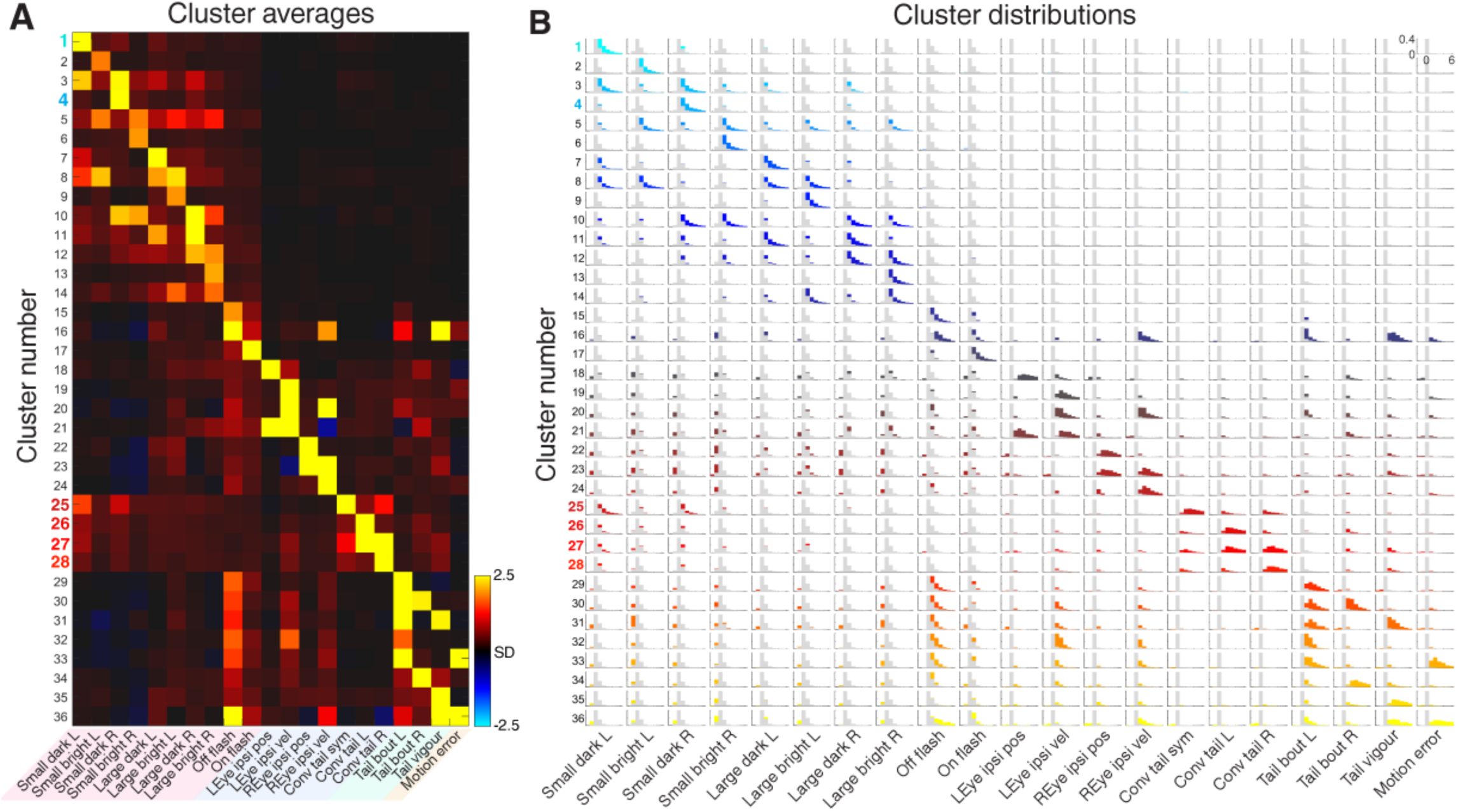
Cluster centroids and distributions. **A** Cluster centroids (mean VMVs) for all 36 clusters. **B** Distributions of VMV components for all clusters. Distributions across all clustered neurons are overlaid in grey. Y-axis ranges from 0 to 0.4 (fraction), x-axis ranges from –2 to 6 (SD).

**Figure 1–supplement 2.**
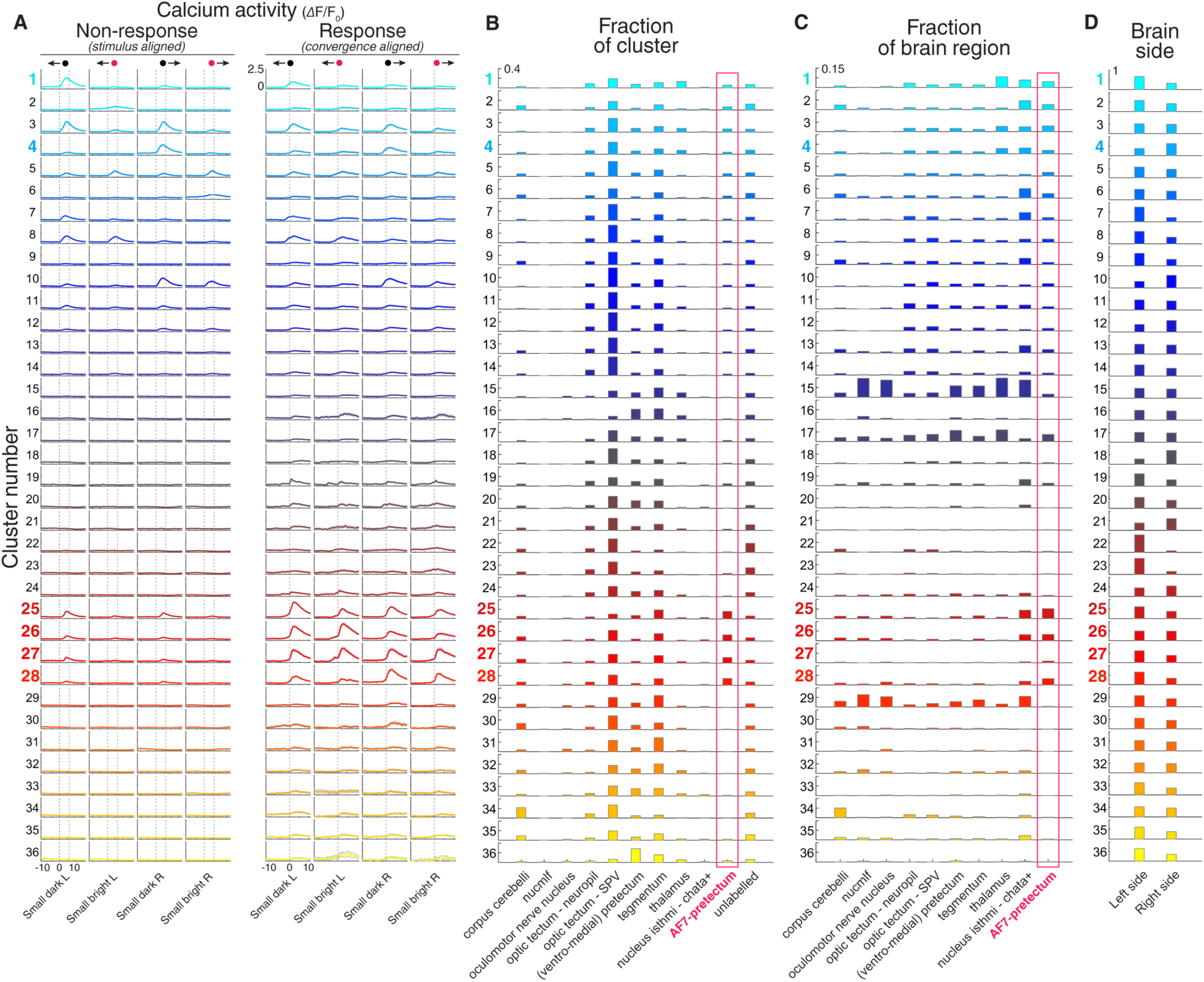
Calcium responses and anatomical distributions of clusters. **A** Visual stimulus-aligned (left four columns) and eye convergence-aligned (right four columns) ΔF/F_0_ responses for all 36 clusters. Responses are shown for small moving spots (dark/bright moving leftwards/rightwards, as indicated at top of columns). Traces show mean ± 95% confidence intervals across all neurons in each cluster. Dashed vertical lines indicate start/end of stimulus presentation in non-response trials (left four columns), or time of eye convergence (right four columns). L, leftwards; R, rightwards. X-axis reports time (s). **B** Anatomical location of clusters (N = 8 larvae). The fraction of cells in each cluster falling within each ZBB anatomical region is shown. Red box highlights AF7-pretectum. Y-axis ranges from 0 to 0.4 (fraction). **C** Fraction of imaged cells within each anatomical region that were assigned to each cluster type. Red box highlights AF7-pretectum. Y-axis ranges from 0 to 0.15 (fraction of imaged neurons in brain region). **D** Fraction of neurons in each cluster located in the left of right brain hemisphere. Y-axis ranges from 0 to 1 (fraction).

**Figure 1–supplement 3.**
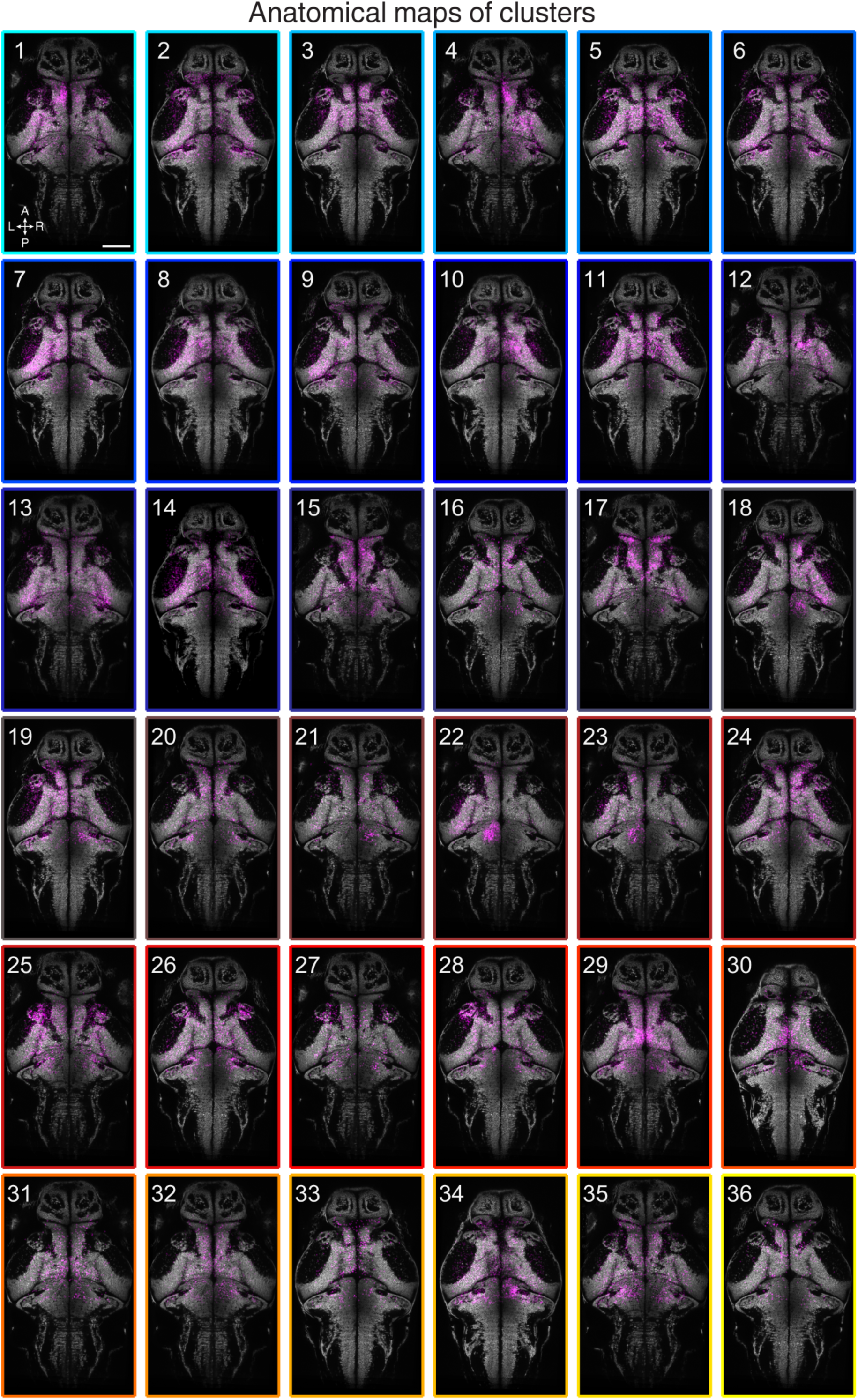
Anatomical maps of clusters. Images show dorsal views of intensity sum projections of all neuronal masks in each cluster (magenta) after registration to the *elavl3:H2B-GCaMP6s* reference brain (grey). Projections (obtained through all focal planes, ∼100 *µ*m total depth) are overlaid on a maximum-intensity projection image (gray) from the *elavl3:H2B-GCaMP6s* reference brain (5 planes, 5 *µ*m depth, from focal planes with the largest number of neurons in each cluster). Scale bar, 100 *µ*m. A, anterior; L, left; P, posterior; R, right.

**Figure 2–supplement 1.**
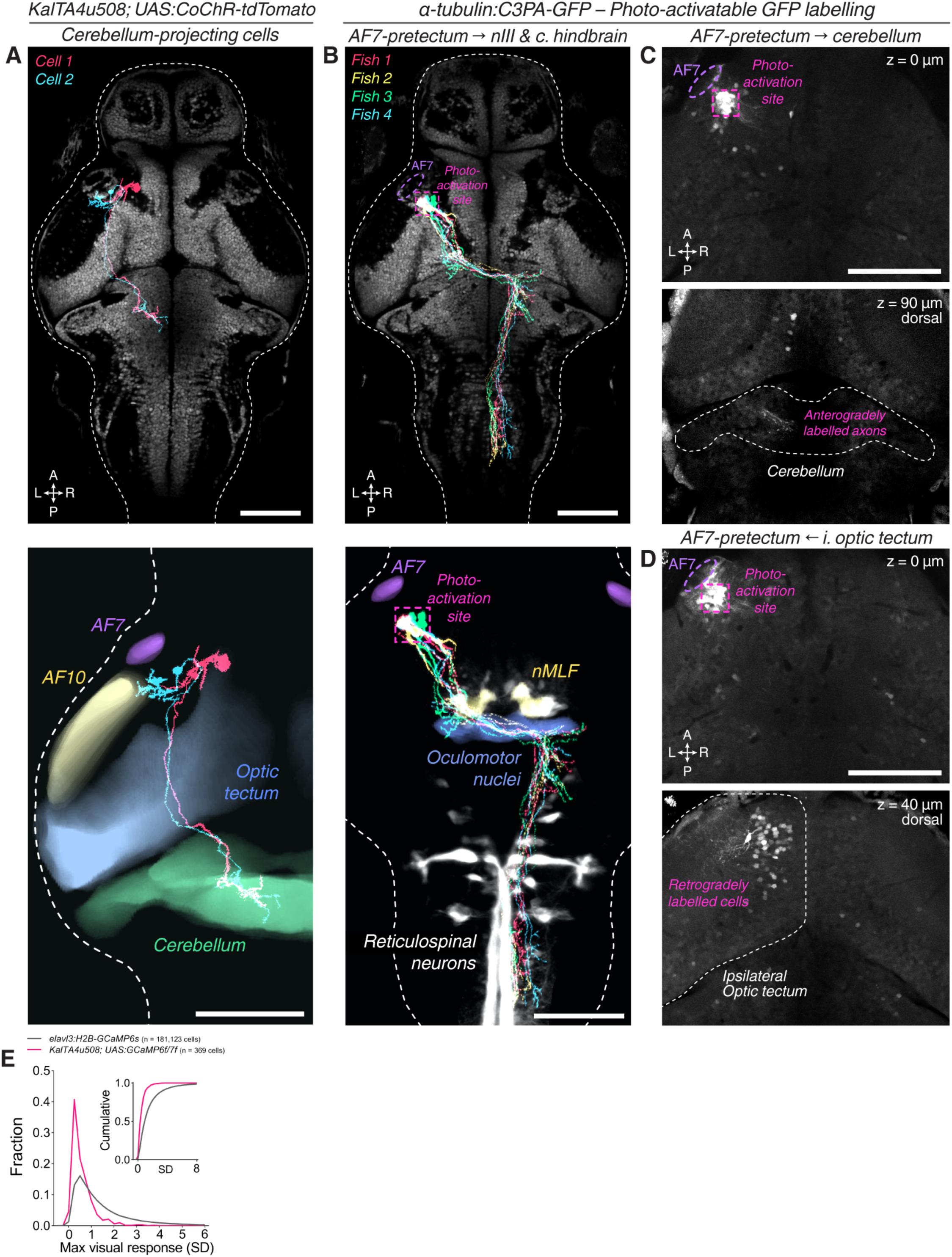
*KalTA4u508* neurons innervating cerebellum, and PA-GFP projection mapping from AF7-pretectum. **A** Tracings *KalTA4u508* neurons projecting to ipsilateral medial corpus cerebellum (n = 2 cells from 2 fish). The bottom image shows tracings overlaid with selected anatomical regions from the ZBB brain atlas. **B** Tracings of PA-GFP-labelled AF7-pretectal cells projecting to oculomotor nuclei and contralateral hindbrain in 7 dpf *α-tubulin:C3PA-GFP* larvae (N = 4 fish) registered to the *elavl3:H2B-GCaMP6s* reference brain (grey). The photo-activation site is indicated in magenta. **C** PA-GFP-labelled AF7-pretectal cells in a 7 dpf *α-tubulin:C3PA-GFP* larva. The photo-activation site is indicated in magenta. Anterogradely labelled axonal terminals are visible in the ipsilateral medial cerebellum (bottom image, z-plane location is relative to top z-plane). **D** A second example of photoactivation that retrogradely labelled cell bodies in the ipsilateral anterior-ventral optic tectum. **E** Distributions of maximum responses across visual stimuli for all recorded neurons in 6–7 dpf *elavl3:H2B-GCaMP6s* larvae (grey, n = 181,123 cells from 8 fish) and *KalTA4u508;UAS:GCaMP6f, or KalTA4u508;UAS:jGCaMP7f* larvae (magenta, n = 369 cells from 30 fish). Before determining the maximum responses for each neuron, mean integrated ΔF/F_0_ for each visual stimulus was normalised by dividing values by the corresponding standard deviation (SD) across all neurons from *elavl3:H2B-GCaMP6s* larvae. Scale bars, 100 *µ*m. A, anterior; c., contralateral; i., ipsilateral; L, left; P, posterior; R, right.

**Figure 3–supplement 1.**
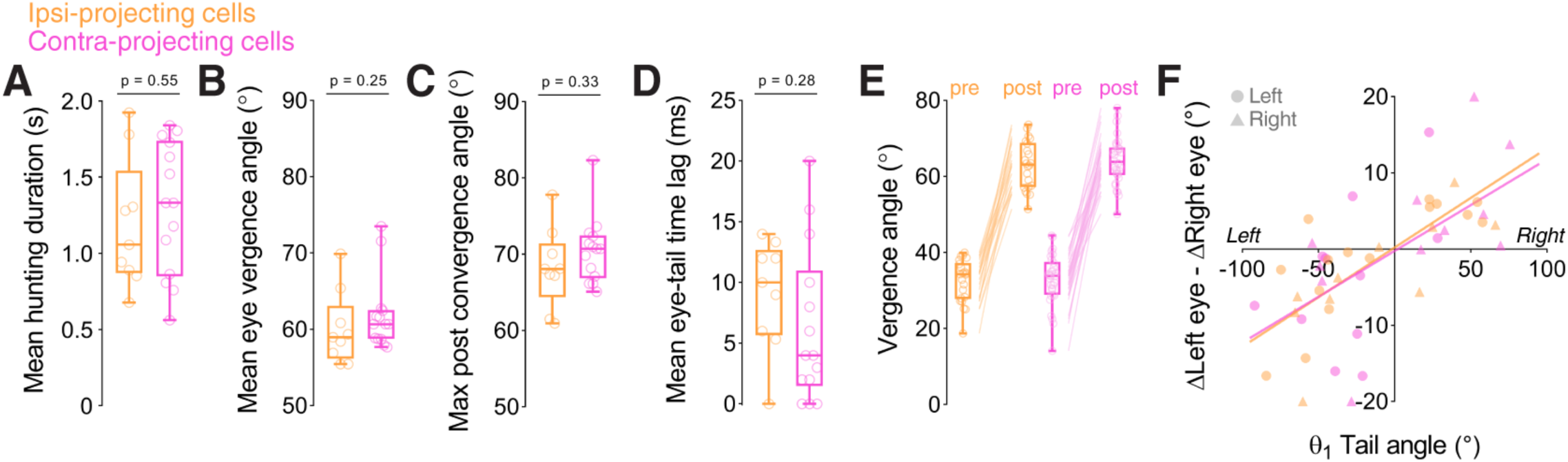
Behavioural kinematics of optogenetically induced hunting. **A–F** Behavioural kinematics for hunting events evoked by optogenetic stimulation of single ipsi-projecting (orange, n = 9 cells) and contra-projecting *KalTA4u508* neurons (magenta, n = 14 cells). In **E** and **F**, data from all hunting events are plotted, whereas in the other plots the mean or maximum for each larva is reported.

**Figure 4–supplement 1.**
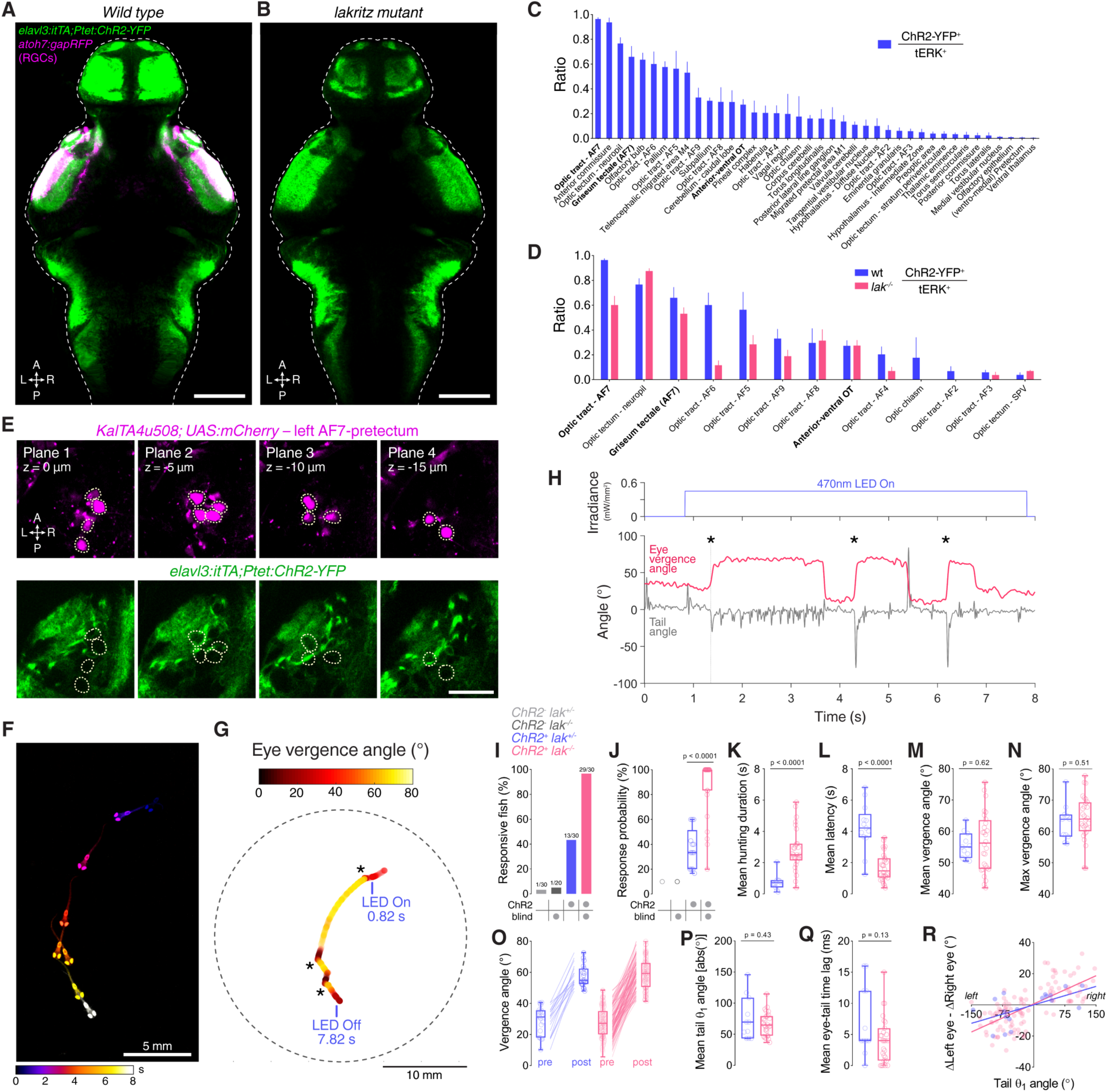
Optogenetic induction of hunting in *elavl3:itTA;Ptet:ChR2-YFP;atoh7:gapRFP* larvae. **A** Dorsal view of ChR2-YFP expression in immunostained 6 dpf *elavl3:itTA;Ptet:ChR2-YFP;atoh7:gapRFP* larvae registered to the ZBB brain atlas. Axonal projections of RGCs labelled by the *atoh7:gapRFP* transgene are displayed in magenta (median of N = 6 fish). Image represents a median of multiple registered single brains (N = 6 fish) and shows a maximum-intensity projection through focal planes encompassing AF7-pretectal regions. **B** ChR2-YFP expression in *elavl3:itTA;Ptet:ChR2-YFP;atoh7:gapRFP;lak*^-/-^ larvae (median of N = 7 fish). Note that no RGC axonal projections are present in the *lakritz* mutant. **C** Overlap between ChR2-YFP and tERK immunostain in *elavl3:itTA;Ptet:ChR2-YFP* larvae, computed as ratio between ChR2-YFP-positive voxels and tERK-positive voxels in each brain region. Mean + SEM values are reported only for anatomical regions showing overlap greater than zero (N = 6 fish). AF7-pretectum encompasses the ZBB masks ‘Optic tract - AF7’ and ‘Griseum tectale (AF7)’. **D** Overlap between ChR2-YFP and tERK immunostain in control (blue) and *lakritz* (pink) larvae (N = 7 fish). **E** ChR2 expression relative to *KalTA4u508* neurons in a 6 dpf *KalTA4u508;UAS:mCherry;elavl3:itTA;Ptet:ChR2-YFP* larva. Images are single focal planes obtained from the left AF7-pretectum (plane 1 is dorsal relative to the other z-planes). *KalTA4u508* neurons are not labelled with ChR2-YFP. **F** Time sequence composite image showing selected frames from an example optogenetically induced hunting sequence in a 6 dpf *elavl3:itTA;Ptet:ChR2-YFP;atoh7:gapRFP;lak*^-/-^ larva (see also Video 3). **G** Vergence angle overlaid onto larval location during the optogenetically induced hunting sequence from **F**. Asterisks indicate time of eye convergences (see also Video 3). **H** Behavioural tracking of tail angle (grey) and ocular vergence angle (red) during the example optogenetically induced hunting sequences shown in **F** and **G**. Asterisks indicate time of eye convergences (see also Video 3). **I** Fraction of larvae that performed eye convergences during optogenetic stimulations. Opsin-positive larvae were either *elavl3:itTA;Ptet:ChR2-YFP;atoh7:gapRFP* or *elavl3:itTA;Ptet:ChR2-YFP;atoh7:gapRFP;lak*^-/-^, whereas opsin-negative larvae were either *atoh7:gapRFP* or *atoh7:gapRFP;lak*^-/-^. Numbers of responsive larvae are reported above the bars. **J** Response probability of larvae that performed eye convergence in at least one optogenetic stimulation trial. **K–R** Behavioural kinematics for optogenetically induced hunting events elicited in sighted (blue, N = 13 larvae) and blind *lak*^-/-^ (pink, N = 29 larvae) *elavl3:itTA;Ptet:ChR2-YFP;atoh7:gapRFP* larvae. In **O** and **R**, data from all hunting events are plotted, whereas in the other plots the mean or maximum for each larva is reported. Scale bars, 100 *µ*m, except in **E**, 30 *µ*m. A, anterior; L, left; P, posterior; R, right.

**Figure 4–supplement 2.**
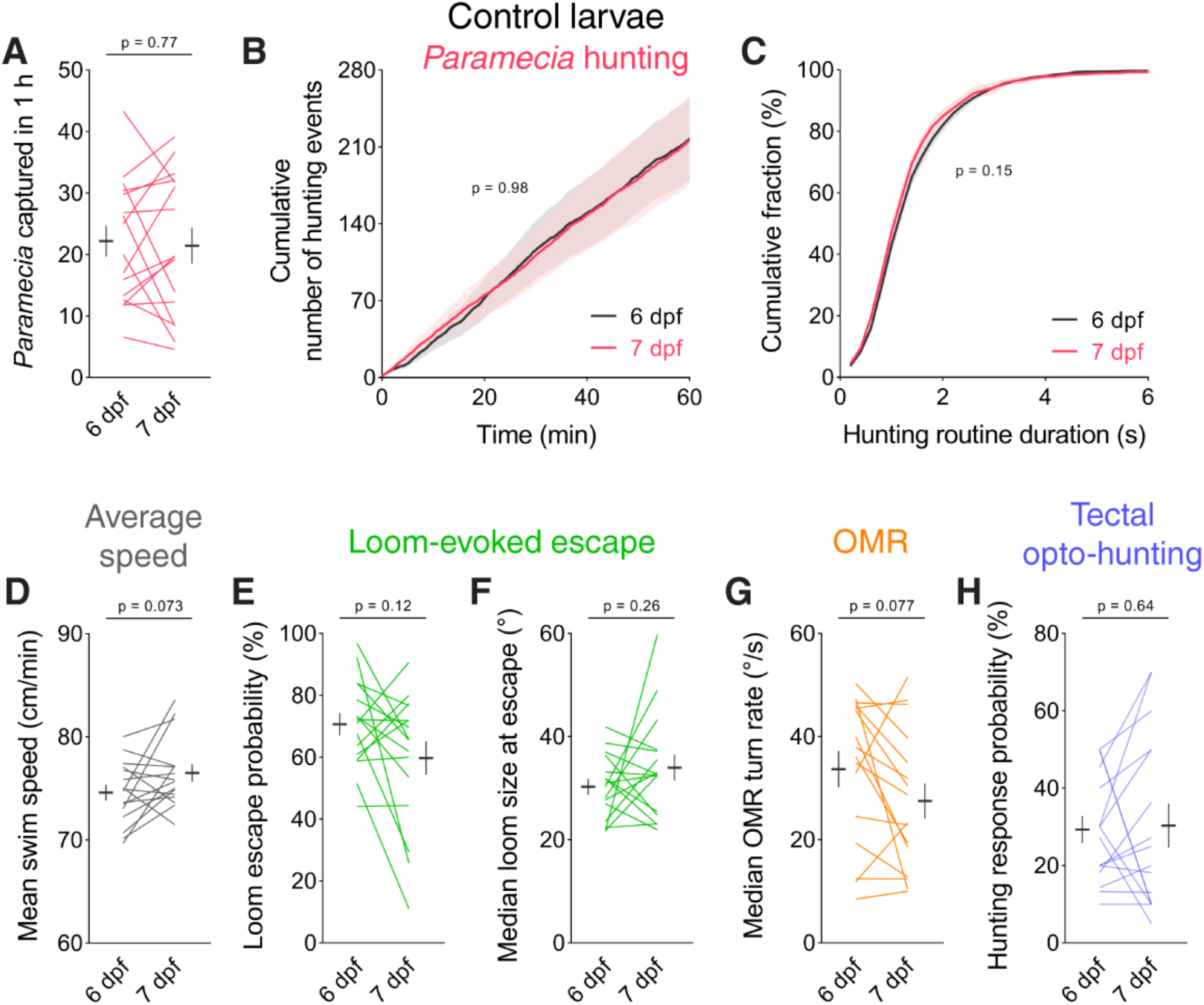
No change in visuomotor behaviours in control larvae. **A–C** Assessment of *Paramecia* hunting in control *KalTA4u508;UAS:mCherry;elavl3:itTA;Ptet:ChR2-YFP* larvae assessed at 6 and 7 dpf (N = 16 larvae). Mean ± SEM is reported for each group. **D** Average swim speed in control larvae. **E–F** Loom-evoked escape behaviour in control larvae. **G** OMR performance in control larvae. **H** Hunting response probability upon optogenetic stimulation of avOT in control larvae.

**Video 1. Z-stack of transgenic line used for calcium imaging with annotated RGC arborisation fields**

Imaging volume (z-stack) of 6 dpf *elavl3:H2B-GCaMP6s;atoh7:gapRFP* brain (mean of N = 3 fish) with labelled RGC arborisation fields (AFs). The green channel shows the *elavl3:H2B-GCaMP6s* reference brain used for anatomical registration.

**Video 2. Hunting behaviour evoked by optogenetic stimulation of a single pretectal neuron.**

Hunting behaviour evoked by optogenetic stimulation of a single ipsi-projecting *KalTA4u508* neuron located in the left AF7-pretectum (this neuron is ‘Cell 4’ in Figure 2F). Tracking data is reported in Figure 3D,E. The video was acquired at 250 frames per second and plays at 0.4 times the original speed. The raw movie is showed on the left, and a background-subtracted inset centred on the larva is showed on the right.

**Video 3. Hunting behaviour evoked by optogenetic stimulation of the anterior-ventral optic tectum.**

Optogenetically induced hunting behaviour in a *elavl3:itTA;Ptet:ChR2-YFP;atoh7:gapRFP;lak*^-/-^ larva. Tracking data is reported in Figure 4–supplement 1F–H. The video was acquired at 250 frames per second and plays at 0.4 times the original speed. The raw movie is showed on the left, and a background-subtracted inset centred on the larva is showed on the right.

## References

Akitake, C.M., Macurak, M., Halpern, M.E., and Goll, M.G. (2011). Transgenerational analysis of transcriptional silencing in zebrafish. Dev Biol 352, 191–201.

Anjum, F., Turni, H., Mulder, P.G., van der Burg, J., and Brecht, M. (2006). Tactile guidance of prey capture in Etruscan shrews. Proc Natl Acad Sci U S A 103, 16544–16549.

Auer, T.O., Duroure, K., De Cian, A., Concordet, J.P., and Del Bene, F. (2014). Highly efficient CRISPR/Cas9-mediated knock-in in zebrafish by homology-independent DNA repair. Genome Res 24, 142–153.

Avants, B.B., Tustison, N.J., Song, G., Cook, P.A., Klein, A., and Gee, J.C. (2011). A reproducible evaluation of ANTs similarity metric performance in brain image registration. Neuroimage 54, 2033–2044.

Bels, V.L., Aerts, P., Chardon, M., Vandewalle, P., Berkhoudt, H., Crompton, A., de Vree, F., Dullemeijer, P., Ewert, J., and Frazzetta, T. (2012). Biomechanics of feeding in vertebrates, Vol 18 (Springer Science & Business Media).

Bianco, I.H., and Engert, F. (2015). Visuomotor transformations underlying hunting behavior in zebrafish. Curr Biol 25, 831–846.

Bianco, I.H., Kampff, A.R., and Engert, F. (2011). Prey capture behavior evoked by simple visual stimuli in larval zebrafish. Front Syst Neurosci 5, 101.

Bianco, I.H., Ma, L.H., Schoppik, D., Robson, D.N., Orger, M.B., Beck, J.C., Li, J.M., Schier, A.F., Engert, F., and Baker, R. (2012). The tangential nucleus controls a gravito-inertial vestibuloocular reflex. Curr Biol 22, 1285–1295.

Borla, M.A., Palecek, B., Budick, S., and O’Malley, D.M. (2002). Prey capture by larval zebrafish: evidence for fine axial motor control. Brain Behav Evol 60, 207–229.

Brainard, D.H. (1997). The Psychophysics Toolbox. Spat Vis 10, 433–436.

Breiman, L., Meisel, W., and Purcell, E.J.T. (1977). Variable kernel estimates of multivariate densities. Technometrics 19, 135–144.

Comoli, E., Ribeiro-Barbosa, E.R., Negrao, N., Goto, M., and Canteras, N.S. (2005). Functional mapping of the prosencephalic systems involved in organizing predatory behavior in rats. Neuroscience 130, 1055–1067.

Dana, H., Sun, Y., Mohar, B., Hulse, B., Hasseman, J.P., Tsegaye, G., Tsang, A., Wong, A., Patel, R., Macklin, J.J., et al. (2018). High-performance GFP-based calcium indicators for imaging activity in neuronal populations and microcompartments. bioRxiv, 434589.

Davison, J.M., Akitake, C.M., Goll, M.G., Rhee, J.M., Gosse, N., Baier, H., Halpern, M.E., Leach, S.D., and Parsons, M.J. (2007). Transactivation from Gal4-VP16 transgenic insertions for tissue-specific cell labeling and ablation in zebrafish. Dev Biol 304, 811–824.

Distel, M., Wullimann, M.F., and Koster, R.W. (2009). Optimized Gal4 genetics for permanent gene expression mapping in zebrafish. Proc Natl Acad Sci U S A 106, 13365–13370.

Dunn, T.W., Gebhardt, C., Naumann, E.A., Riegler, C., Ahrens, M.B., Engert, F., and Del Bene, F. (2016). Neural Circuits Underlying Visually Evoked Escapes in Larval Zebrafish. Neuron 89, 613–628.

Ewert, J.P. (1970). Neural mechanisms of prey-catching and avoidance behavior in the toad (Bufo bufo L.). Brain Behav Evol 3, 36–56.

Ewert, J.P. (1997). Neural correlates of key stimulus and releasing mechanism: a case study and two concepts. Trends Neurosci 20, 332–339.

Ewert, J.P., Buxbaum-Conradi, H., Dreisvogt, F., Glagow, M., Merkel-Harff, C., Rottgen, A., Schurg-Pfeiffer, E., and Schwippert, W.W. (2001). Neural modulation of visuomotor functions underlying prey-catching behaviour in anurans: perception, attention, motor performance, learning. Comp Biochem Physiol A Mol Integr Physiol 128, 417–461.

Fajardo, O., Zhu, P., and Friedrich, R.W. (2013). Control of a specific motor program by a small brain area in zebrafish. Front Neural Circuits 7, 67.

Flood, T.F., Iguchi, S., Gorczyca, M., White, B., Ito, K., and Yoshihara, M. (2013). A single pair of interneurons commands the Drosophila feeding motor program. Nature 499, 83–87.

Frost, W.N., and Katz, P.S. (1996). Single neuron control over a complex motor program. Proc Natl Acad Sci U S A 93, 422–426.

Gahtan, E., Tanger, P., and Baier, H. (2005). Visual prey capture in larval zebrafish is controlled by identified reticulospinal neurons downstream of the tectum. J Neurosci 25, 9294–9303.

Han, W., Tellez, L.A., Rangel, M.J., Jr., Motta, S.C., Zhang, X., Perez, I.O., Canteras, N.S., Shammah-Lagnado, S.J., van den Pol, A.N., and de Araujo, I.E. (2017). Integrated Control of Predatory Hunting by the Central Nucleus of the Amygdala. Cell 168, 311–324.e318.

Helmbrecht, T.O., Dal Maschio, M., Donovan, J.C., Koutsouli, S., and Baier, H. (2018). Topography of a Visuomotor Transformation. Neuron 100, 1429–1445.e1424.

Herrero, L., Rodriguez, F., Salas, C., and Torres, B. (1998). Tail and eye movements evoked by electrical microstimulation of the optic tectum in goldfish. Exp Brain Res 120, 291–305.

Horstick, E.J., Jordan, D.C., Bergeron, S.A., Tabor, K.M., Serpe, M., Feldman, B., and Burgess, H.A. (2015). Increased functional protein expression using nucleotide sequence features enriched in highly expressed genes in zebrafish. Nucleic Acids Res 43, e48.

Jing, J. (2009). Command systems. In Encyclopedia of neuroscience (Elsevier Ltd), pp. 1149–1158.

Jordi, J., Guggiana-Nilo, D., Soucy, E., Song, E.Y., Lei Wee, C., and Engert, F. (2015). A high-throughput assay for quantifying appetite and digestive dynamics. Am J Physiol Regul Integr Comp Physiol 309, R345–357.

Kawashima, T., Zwart, M.F., Yang, C.T., Mensh, B.D., and Ahrens, M.B. (2016). The Serotonergic System Tracks the Outcomes of Actions to Mediate Short-Term Motor Learning. Cell 167, 933–946.e920.

Kay, J.N., Finger-Baier, K.C., Roeser, T., Staub, W., and Baier, H. (2001). Retinal ganglion cell genesis requires lakritz, a Zebrafish atonal Homolog. Neuron 30, 725–736.

Klapoetke, N.C., Murata, Y., Kim, S.S., Pulver, S.R., Birdsey-Benson, A., Cho, Y.K., Morimoto, T.K., Chuong, A.S., Carpenter, E.J., Tian, Z., et al. (2014). Independent optical excitation of distinct neural populations. Nat Methods 11, 338–346.

Knafo, S., Fidelin, K., Prendergast, A., Tseng, P.-E.B., Parrin, A., Dickey, C., Böhm, U.L., Figueiredo, S.N., Thouvenin, O., Pascal-Moussellard, H., et al. (2017). Mechanosensory neurons control the timing of spinal microcircuit selection during locomotion. Elife 6, e25260.

Kohl, J., Babayan, B.M., Rubinstein, N.D., Autry, A.E., Marin-Rodriguez, B., Kapoor, V., Miyamishi, K., Zweifel, L.S., Luo, L., Uchida, N., et al. (2018). Functional circuit architecture underlying parental behaviour. Nature 556, 326–331.

Korn, H., and Faber, D.S. (2005). The Mauthner cell half a century later: a neurobiological model for decision-making? Neuron 47, 13–28.

Koyama, M., Kinkhabwala, A., Satou, C., Higashijima, S., and Fetcho, J. (2011). Mapping a sensory-motor network onto a structural and functional ground plan in the hindbrain. Proc Natl Acad Sci U S A 108, 1170–1175.

Kupfermann, I., and Weiss, K.R. (1978). The command neuron concept. Behavioural and Brain Sciences 1, 3–39.

Li, Y., Zeng, J., Zhang, J., Yue, C., Zhong, W., Liu, Z., Feng, Q., and Luo, M. (2018). Hypothalamic Circuits for Predation and Evasion. Neuron 97, 911–924.e915.

Lister, J.A., Robertson, C.P., Lepage, T., Johnson, S.L., and Raible, D.W. (1999). nacre encodes a zebrafish microphthalmia-related protein that regulates neural-crest-derived pigment cell fate. Development 126, 3757–3767.

Longair, M.H., Baker, D.A., and Armstrong, J.D. (2011). Simple Neurite Tracer: open source software for reconstruction, visualization and analysis of neuronal processes. Bioinformatics 27, 2453–2454.

Marquart, G.D., Tabor, K.M., Brown, M., Strykowski, J.L., Varshney, G.K., LaFave, M.C., Mueller, T., Burgess, S.M., Higashijima, S., and Burgess, H.A. (2015). A 3D Searchable Database of Transgenic Zebrafish Gal4 and Cre Lines for Functional Neuroanatomy Studies. Front Neural Circuits 9, 78.

Marquart, G.D., Tabor, K.M., Horstick, E.J., Brown, M., Geoca, A.K., Polys, N.F., Nogare, D.D., and Burgess, H.A. (2017). High-precision registration between zebrafish brain atlases using symmetric diffeomorphic normalization. Gigascience 6, 1–15.

Marques, J.C., Lackner, S., Felix, R., and Orger, M.B. (2018). Structure of the Zebrafish Locomotor Repertoire Revealed with Unsupervised Behavioral Clustering. Curr Biol 28, 181–195.e185.

Masai, I., Lele, Z., Yamaguchi, M., Komori, A., Nakata, A., Nishiwaki, Y., Wada, H., Tanaka, H., Nojima, Y., Hammerschmidt, M., et al. (2003). N-cadherin mediates retinal lamination, maintenance of forebrain compartments and patterning of retinal neurites. Development 130, 2479–2494.

McElligott, M.B., and O’Malley D M. (2005). Prey tracking by larval zebrafish: axial kinematics and visual control. Brain Behav Evol 66, 177–196.

Miri, A., Daie, K., Burdine, R.D., Aksay, E., and Tank, D.W. (2011). Regression-based identification of behavior-encoding neurons during large-scale optical imaging of neural activity at cellular resolution. J Neurophysiol 105, 964–980.

Muto, A., Lal, P., Ailani, D., Abe, G., Itoh, M., and Kawakami, K. (2017). Activation of the hypothalamic feeding centre upon visual prey detection. Nat Commun 8, 15029.

Park, S.G., Jeong, Y.C., Kim, D.G., Lee, M.H., Shin, A., Park, G., Ryoo, J., Hong, J., Bae, S., Kim, C.H., et al. (2018). Medial preoptic circuit induces hunting-like actions to target objects and prey. Nat Neurosci 21, 364–372.

Patterson, B.W., Abraham, A.O., MacIver, M.A., and McLean, D.L. (2013). Visually guided gradation of prey capture movements in larval zebrafish. J Exp Biol 216, 3071–3083.

Qian, J., Hastie, T., Friedman, J., Tibshirani, R., and Simon, N. (2013). Glmnet for matlab. available at: http://www.stanford.edu/~hastie/glmnet_matlab/.

Semmelhack, J.L., Donovan, J.C., Thiele, T.R., Kuehn, E., Laurell, E., and Baier, H. (2014). A dedicated visual pathway for prey detection in larval zebrafish. Elife 3.

Sillar, K.T., Picton, L., and Heitler, W.J. (2016). The neuroethology of predation and escape (Wiley Blackwell).

Trivedi, C.A., and Bollmann, J.H. (2013). Visually driven chaining of elementary swim patterns into a goal-directed motor sequence: a virtual reality study of zebrafish prey capture. Front Neural Circuits 7, 86.

Vladimirov, N., Mu, Y., Kawashima, T., Bennett, D.V., Yang, C.T., Looger, L.L., Keller, P.J., Freeman, J., and Ahrens, M.B. (2014). Light-sheet functional imaging in fictively behaving zebrafish. Nat Methods 11, 883–884.

von Philipsborn, A.C., Liu, T., Yu, J.Y., Masser, C., Bidaye, S.S., and Dickson, B.J. (2011). Neuronal control of Drosophila courtship song. Neuron 69, 509–522.

Yanez, J., Suarez, T., Quelle, A., Folgueira, M., and Anadon, R. (2018). Neural connections of the pretectum in zebrafish (Danio rerio). J Comp Neurol 526, 1017–1040.

Yoshihara, M., and Yoshihara, M. (2018). ‘Necessary and sufficient’ in biology is not necessarily necessary - confusions and erroneous conclusions resulting from misapplied logic in the field of biology, especially neuroscience. J Neurogenet 32, 53–64.

Zolessi, F.R., Poggi, L., Wilkinson, C.J., Chien, C.B., and Harris, W.A. (2006). Polarization and orientation of retinal ganglion cells in vivo. Neural Dev 1, 2.

Zou, H., and Hastie, T. (2005). Regularization and variable selection via the elastic net. 67, 301–320.

